# Structural and functional dissection of the Pacinian corpuscle reveals an active role of the inner core in touch detection

**DOI:** 10.1101/2024.08.24.609509

**Authors:** Luke H. Ziolkowski, Yury A. Nikolaev, Akitoshi Chikamoto, Mai Oda, Viktor V. Feketa, David Monedero-Alonso, Serafima A. Ardasheva, Samuel S. Bae, C. Shan Xu, Song Pang, Elena O. Gracheva, Sviatoslav N. Bagriantsev

## Abstract

Pacinian corpuscles are rapidly adapting mechanoreceptor end-organs that detect transient touch and high-frequency vibration. In the prevailing model, these properties are determined by the outer core, which acts as a mechanical filter limiting static and low-frequency stimuli from reaching the afferent terminal—the sole site of touch detection in corpuscles. Here, we determine the detailed 3D architecture of corpuscular components and reveal their contribution to touch detection. We show that the outer core is dispensable for rapid adaptation and frequency tuning. Instead, these properties arise from the inner core, composed of gap junction-coupled lamellar Schwann cells (LSCs) surrounding the afferent terminal. By acting as additional touch sensing structures, LSCs potentiate mechanosensitivity of the terminal, which detects touch via fast-inactivating ion channels. We propose a model in which Pacinian corpuscle function is mediated by an interplay between mechanosensitive LSCs and the afferent terminal in the inner core.

**Highlights:**

- eFIB-SEM reveals detailed 3D architecture of the entire Pacinian (Herbst) corpuscle
- Inner, not outer core mediates rapid adaptation and frequency tuning
- Afferent terminal detects touch via fast-inactivating ion channels
- Mechanosensitive lamellar Schwann cells tune afferent terminal sensitivity to touch

## Introduction

The sense of mechanical touch is indispensable for everyday life, enabling interaction with the physical world, detection of pain and pleasure, formation of social bonds, and manipulation of tools and objects. Mammalian Pacinian corpuscles and their avian homologs (historically known as Herbst corpuscles, and herein referred to as avian Pacinians) are located in the skin and periosteum, where they detect transient touch and high-frequency vibration (Cobo et al., 2021; Gottschaldt, 1974; Handler and Ginty, 2021; Lee et al., 2024; Schneider et al., 2016; Talbot et al., 1968; Turecek and Ginty, 2024; Ziolkowski et al., 2022). These properties stem from the ability of corpuscles to respond only to dynamic, but not static stimuli (a process called rapid adaptation) and exhibit increased sensitivity to high-frequency vibration (high-pass frequency filtering). Despite the variation in size and anatomical location, Pacinian corpuscles from different species exhibit comparable overall architecture and sensory properties (Bell et al., 1994; Bolanowski et al., 1994; Bolanowski and Zwislocki, 1984; Dorward and McIntyre, 1971; Handler et al., 2023; Saxod, 1996; Zelena et al., 1997; Ziolkowski *et al*., 2022), suggesting a unifying mechanism of touch detection, which remains obscure.

The exterior capsule of Pacinian corpuscles, known as the outer core, is formed by several layers of outer core lamellar cells (OCLCs). The outer core forms a diffusion-restricting barrier that maintains a turgor pressure around the inner core, which is composed of lamellar Schwann cells (LSCs) surrounding the terminal of a mechanoreceptor afferent (Bell *et al*., 1994). The currently accepted view is that the multilayered structure of the outer core together with turgor pressure in the inner cavity form a mechanical filter restricting static and low-frequency mechanical stimuli from reaching the afferent terminal (Loewenstein and Skalak, 1966; Quindlen et al., 2016; Suazo et al., 2022). Thus, the outer core has been proposed as the main structural component enabling rapid adaptation and frequency filtering in Pacinian corpuscles. However, experimental and modeling data show that these properties remain largely invariant even though the number of outer core layers varies from a few to several hundred among different species, and often increases over the organism’s lifetime (Quindlen-Hotek et al., 2020). These observations call for further investigation of the role of the outer core and other structural components in shaping the functional tuning of Pacinian corpuscles.

In the inner core, each LSC sprouts numerous semi-concentric processes, known as inner core lamellae, around the afferent terminal. While the terminal is thought to be the sole site of touch detection in Pacinian corpuscles, LSCs were hypothesized to play structural, developmental, supportive, and trophic roles (Loewenstein and Mendelson, 1965; Logan et al., 2024; Meltzer et al., 2022; Mendelson and Loewenstein, 1964; Pawson et al., 2009; Suazo *et al*., 2022), but whether LSCs actively participate in touch detection is unknown. Additionally, to our knowledge, direct electrophysiological measurements from LSCs or the terminal via patch-clamp recording have not been reported in any species. As a result, biophysical properties of mechanically gated ion channels that perform the mechano-electric conversion at the physiological site of touch detection in Pacinian corpuscles remain unexplored. Understanding the mechanism of Pacinian corpuscle function also requires establishing of the spatial arrangement of its cellular components, including the outer and inner cores, which is currently missing except for the structure of the afferent terminal (Handler *et al*., 2023). In this study, we combined structural and functional studies to determine the 3D architecture of the avian Pacinian corpuscle and reveal the contribution of its cellular components to touch detection.

## Results

### 3D architecture of an avian Pacinian corpuscle

We used enhanced focused ion beam scanning electron microscopy (eFIB-SEM) to determine the detailed architecture of the Pacinian corpuscle in the bill skin of the late-stage embryonic tactile specialist Mallard duck at a voxel size of 8 x 8 x 8 nm (Figure 1A-D, Table S1). Ducks are precocial birds, whose development, including the somatosensory system, reaches a near-complete stage before hatching (Nikolaev et al., 2023; Saxod, 1978; Ziolkowski *et al*., 2022; Ziolkowski et al., 2023). The reconstructed corpuscle had an ovoid shape measuring 66 μm along the long axis, and a diameter in the widest region of 53 μm (Movie S1). Flattened OCLCs form the outer core of the corpuscle (Figure 1C-E), creating a diffusion barrier and a unique ionic environment around the inner core (Bell *et al*., 1994; Berkhoudt, 1980; Gray and Sato, 1955; Ilyinsky et al., 1976). The space between the outer and inner cores is filled with loosely packed collagen fibers (Figure S1). The inner core is composed of a single unbranched mechanoreceptor afferent terminal surrounded by 12 LSCs. The bodies of LSCs containing nuclei are arranged in two columns on opposite sides of the afferent (Figure 1F). Each LSC sprouts numerous thin concentric lamellae with surface area ranging from 10 μm^2^ to 800 μm^2^ that envelop a portion of the afferent terminal (Figure 1G, Figure S2). These inner core lamellae extend from the soma along the length of the terminal and interleave with the lamellae from neighboring LSCs from the same column and from the opposite side of the terminal (Figure 2A).

**Figure 1.**
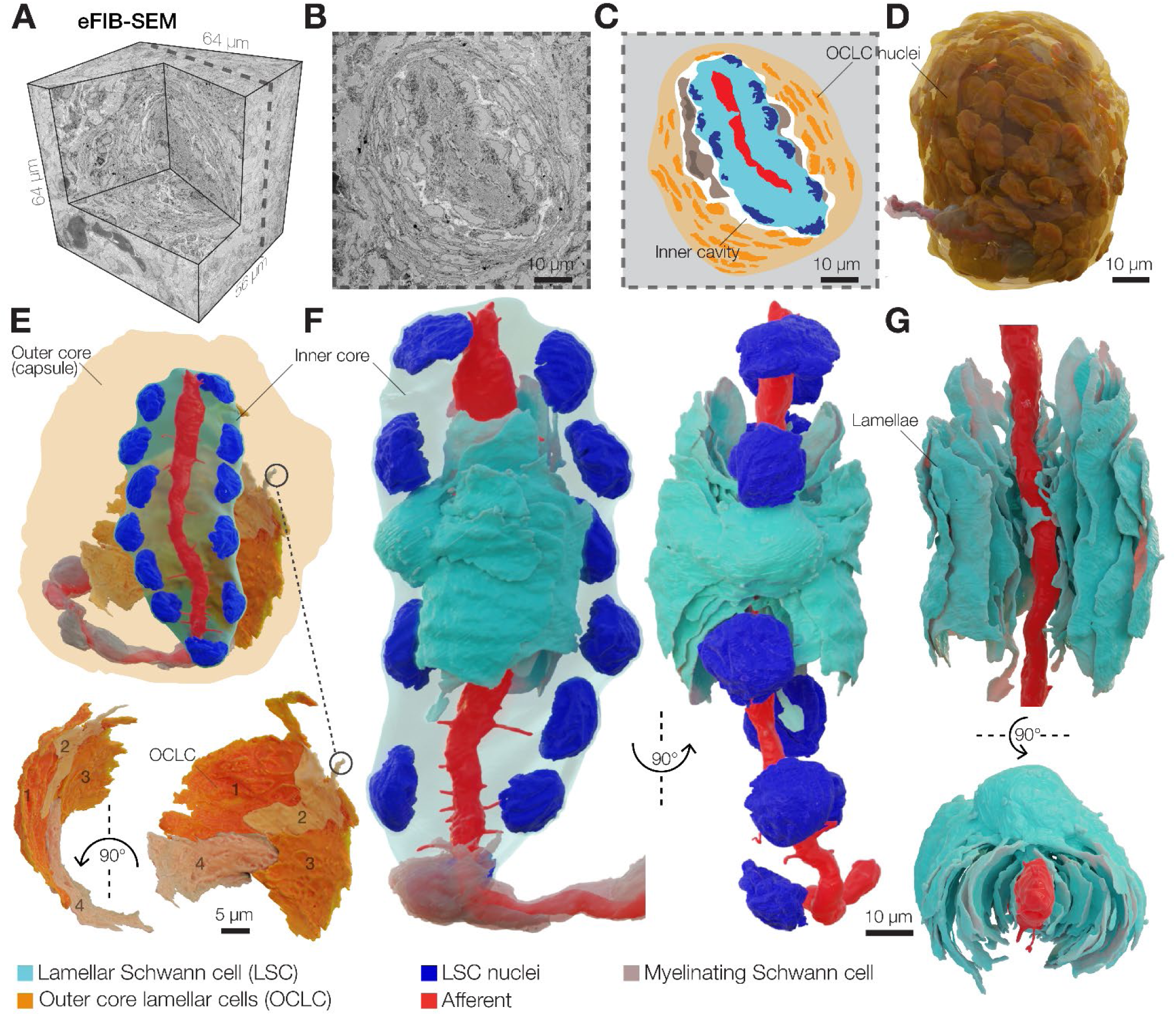
3D architecture of the Pacinian corpuscle. (A) A 3D volume of duck bill skin dermis obtained by eFIB-SEM with 8 nm^3^ resolution. (B, C) A single eFIB-SEM image (B) and an illustration (C) of a section of an avian Pacinian corpuscle. (D) 3D reconstruction of the avian Pacinian corpuscle. (E) 3D reconstruction of the Pacinian corpuscle showing the location of the inner core inside the outer core (top), and reconstruction of four outer core lamellar cells (bottom). (F, G) 3D reconstruction of the inner core showing the architecture of the afferent terminal and one of 12 lamellar Schwann cells (LSC). Different shades of cyan denote lamellae from the same LSC.

**Figure 2.**
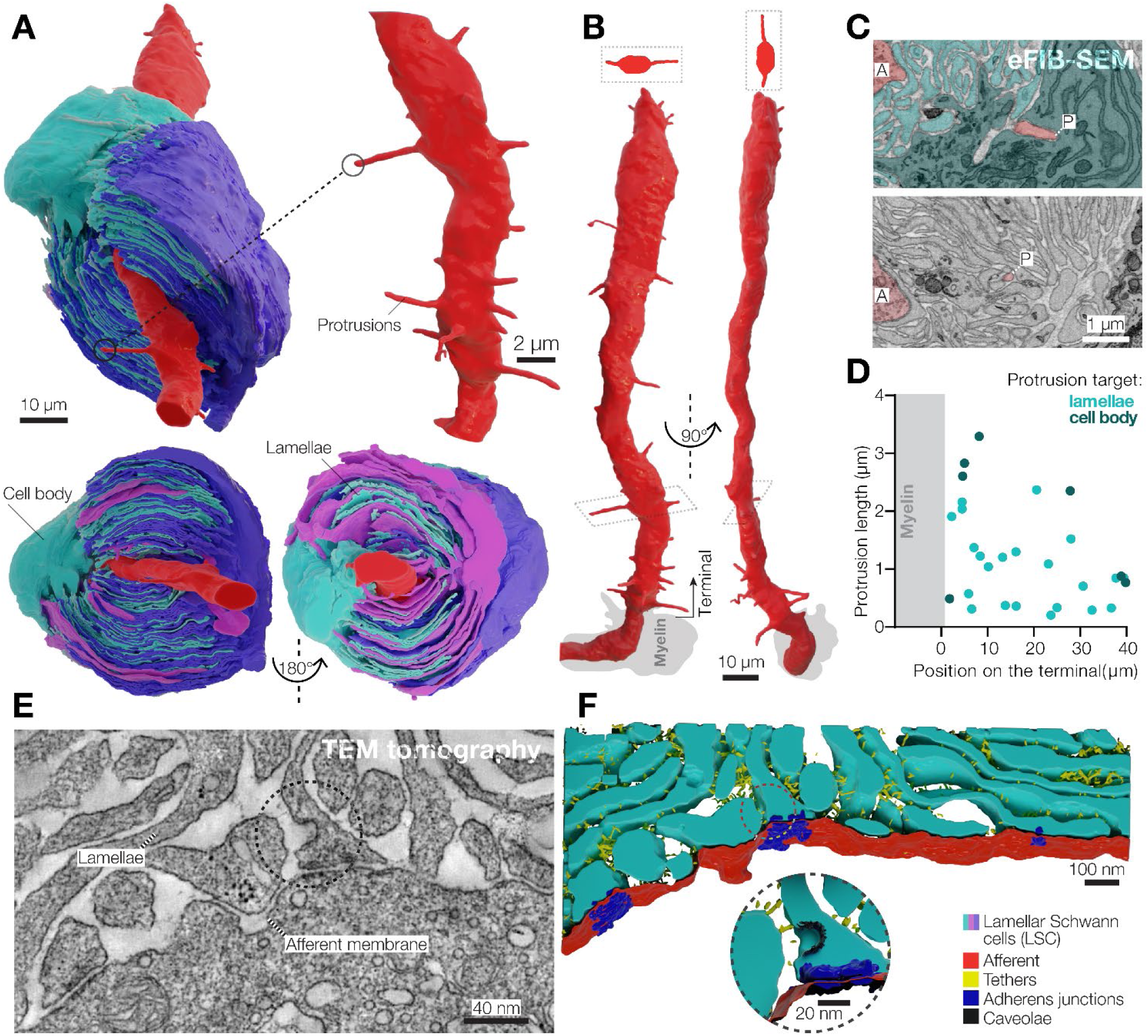
3D architecture of LSCs and the afferent terminal in the Pacinian corpuscle. (A) 3D reconstruction of a pair of opposing LSCs from the inner core (upper panel). An additional LSC is located on top of the cyan LSC (bottom panel). (B) 3D reconstruction of the afferent terminal with 29 protrusions. Two cross section views are shown at the top of the terminal. (C) A pseudo-colored eFIB-SEM image showing protrusion tips targeting the LSC body (upper panel) and lamellae. A, afferent terminal; P, protrusion tip. (D) Localization, length and target of afferent protrusions. (E, F) Transmission electron microscopy image (E) and its 3D reconstruction of the lamellae-afferent contact area.

The corpuscle is innervated by a single mechanoreceptor afferent surrounded by myelinating Schwann cells (Figure 1F). Inside the corpuscle the afferent loses myelination and forms an unbranched 48 μm long terminal that extends through the entire length of the sensory core. When viewed in cross-section, the terminal is elliptical, with its long axis aligned with the cleft in the surrounding inner core lamellae formed by LSCs (Figure S3). The cytosol of the afferent terminal contains numerous densely packed elongated mitochondria (Figure S3A, B). We also detected clear vesicles and occasional dense core vesicles, which were more abundant in the ultra-terminal end compared to the rest of the terminal (Figure S3C, D).

A salient feature of the afferent terminal is its prominent spike-like protrusions, which mostly emanate from the opposing sides of the ovoid afferent and face the apparent cleft in the inner core lamellae (Figure 2A, B and Figure S2). The protrusions, which are considered as putative sites of mechanotransduction (Bolanowski *et al*., 1994; Handler *et al*., 2023; Zelena *et al*., 1997), are located along the entire length of the afferent, have a diameter of ∼250 nm and a length of 0.2-3.3 μm. The tips of most protrusions reached the lamellae of surrounding LSCs, but the longest protrusions extended to the nucleus-containing LSC ‘body’ (Figure 2C, D). We detected 29 protrusions along the terminal, with an average density of 0.6 protrusions per micrometer of terminal length. We also performed eFIB-SEM imaging of a second Pacinian corpuscle. This structure measured 129 μm along the long axis, but had the same overall architecture, including the outer and inner cores, and the afferent terminal (Figure S5A-C, Table S1). The inner core of the second Pacinian had 17 LSCs surrounding a 69 μm long afferent terminal with 0.2-4.5 μm long protrusions and a density of 0.8 protrusions per micrometer of terminal length (Figure S5D, E). In this corpuscle, the terminal ended with an enlarged structure, referred to as the ‘bulb’, which contained clear and dense core vesicles, similar to the first Pacinian (Figure S5D, F). The density of protrusions in the two duck afferent terminals is smaller than in murine Pacinian corpuscles reported elsewhere (Handler *et al*., 2023) and in the accompanying study (Chen et al.) and may be either species-specific, or reflect the developmental stage.

We used 3D transmission electron microscopy tomography to reconstruct and segment a 1.6 x 1.3 x 0.15 μm volume containing inner core lamellae near the afferent terminal of the first corpuscle (Figure 2E). Inner core lamellae contained caveolae and formed contacts with adjacent lamellae via gap junctions (Figure S6), adherens junctions and tethers (Figure 2F). Notably, numerous adherens junctions and tethers also connected inner core lamellae and afferent terminal membranes, demonstrating tight physical coupling between the Schwann and neuronal components of the inner core (Figure 2F and Movie S2).

### Integrity of the outer core is dispensable for frequency tuning and firing adaptation

Pacinian corpuscles in mammals and birds detect transient touch and high-frequency vibration (Bell *et al*., 1994; Dorward and McIntyre, 1971; Gottschaldt, 1974; Handler and Ginty, 2021; Lee *et al*., 2024; Ziolkowski *et al*., 2022). Two characteristic features of corpuscles determine these functions: the cessation of actional potential (AP) firing in the afferent during static stimulation (rapid adaptation), and the decrease of the firing threshold upon repetitive stimulation at high frequency (high-pass frequency tuning). Earlier studies suggested that both these properties are enabled by the multilayered structure of the outer core, which maintains turgor pressure around the inner core and thus acts as a mechanical filter that prevents static and low-frequency stimuli from reaching the afferent terminal. The integrity of the outer core is thus thought to be essential for Pacinian corpuscle function (Bell *et al*., 1994; Gray and Sato, 1955; Loewenstein and Mendelson, 1965; Loewenstein and Skalak, 1966; Mendelson and Loewenstein, 1964; Pease and Quilliam, 1957).

To directly test this hypothesis, we used a previously developed *ex vivo* preparation from late-stage embryonic duck bill skin that enables direct electrophysiological access to corpuscles (Nikolaev et al., 2020; Ziolkowski *et al*., 2023). As the first step, we probed the functional characteristics of intact duck Pacinian corpuscles using single-fiber afferent recordings outside the corpuscle (Figure 3A, B). Step mechanical indentation applied onto the corpuscle evoked APs in the rapidly adapting fashion during the dynamic, but not the static, phases of the stimulus. Further, the APs were inhibited by the voltage-gated sodium channel blocker tetrodotoxin (Figure 3C). In 68.4% of the recordings, firing occurred during both the ON and OFF dynamic phases (Figure 3D). To test whether the integrity of the outer core and the turgor pressure inside the corpuscle are important for rapid adaptation, we compared corpuscle function before and after rupturing the outer core via the combination of mechanical disruption and a high-pressure stream of Krebs solution from a patch pipette (Figure 3E and Methods). Strikingly, this procedure failed to affect the rapid adaptation, as the afferent continued to fire only during the ON and OFF phases and with unchanged threshold of activation (Figure 3F). Next, we assayed the importance of outer core integrity on frequency tuning. Repetitive mechanical stimulation of intact corpuscles with increasing amplitude at different frequencies evoked characteristic high-pass filtering firing in the afferent with peak sensitivity around 100-200Hz. We found that rupturing the outer core also failed to affect frequency tuning (Figure 3G, H). Thus, our data show that, first, late-stage embryonic duck Pacinian corpuscles exhibit rapid adaptation and high-pass frequency filtering similar to corpuscles from adult birds and are thus functionally mature (Dorward and McIntyre, 1971; Gottschaldt, 1974). Second, contrary to the accepted model, the integrity of the outer core and the turgor pressure are dispensable for rapid adaptation and frequency tuning of duck Pacinian corpuscles.

**Figure 3.**
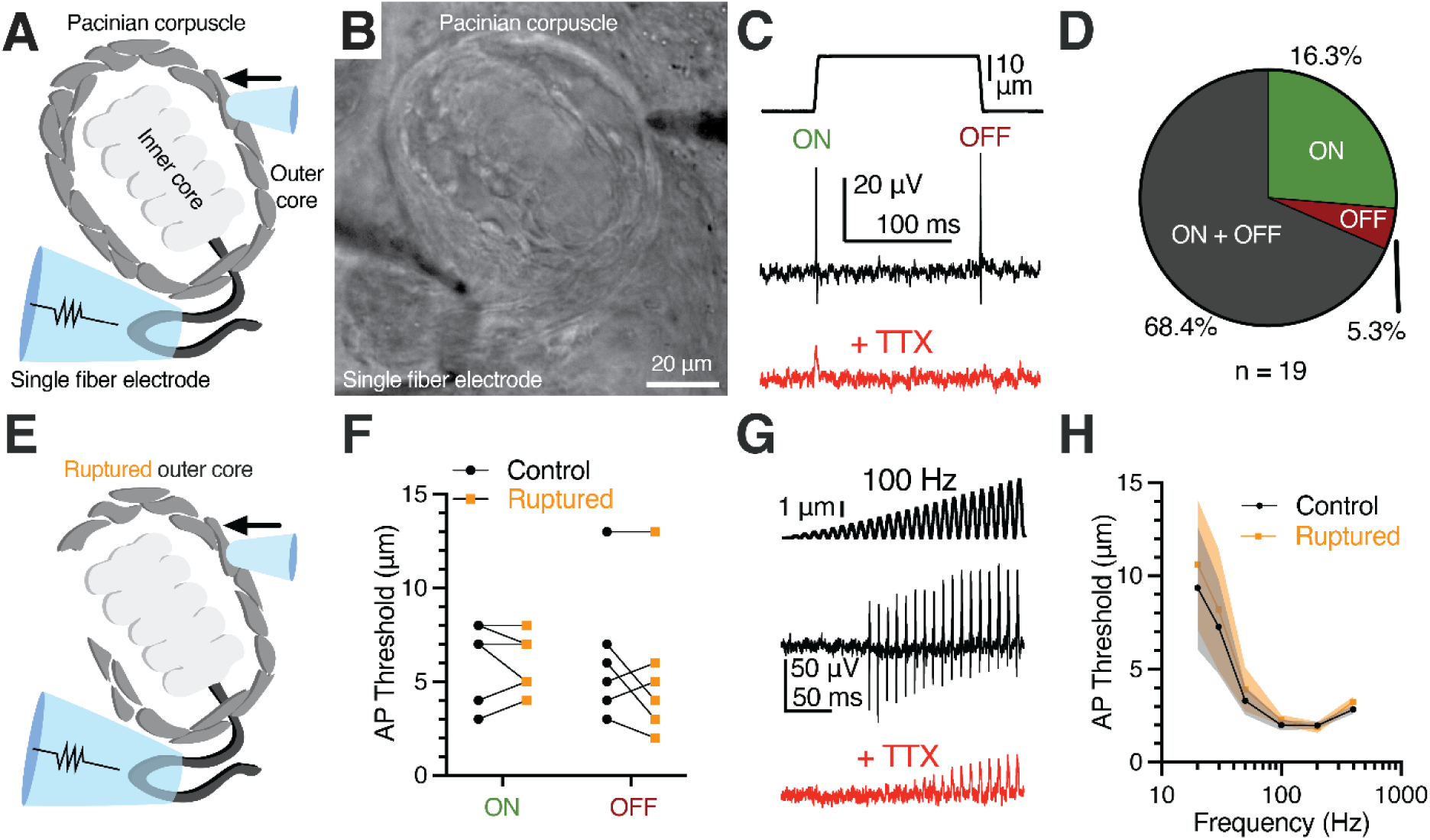
Outer core integrity is dispensable for rapid adaptation and frequency tuning of the Pacinian corpuscle. (A, B) Illustration (A) and bright-field image (B) of Pacinian single-fiber recording. (C) A mechanical step stimulus (top) and a representative single-fiber recording from the Pacinian afferent (middle) with 1 μM TTX added to block APs (bottom). (D) Proportion of Pacinian mechanoreceptors that fire an AP in the dynamic onset (ON), offset (OFF), or both ON and OFF phases. (E) Illustration of a Pacinian corpuscle with the outer layers of OCLCs ruptured. (F) Threshold comparison of the ON and OFF responses using the protocol in (C). Connected symbols represent paired observations from the same corpuscle. No difference between the intact (control) and ruptured outer core conditions was detected (two-way repeated measures ANOVA, p=0.2865). (G) A 100 Hz sinusoidal-ramp mechanical stimulus applied to corpuscles (top) and an exemplar single-fiber response (middle) with 1 μM TTX added to block APs (bottom). (H) Population tuning curve of Pacinian afferents using a sinusoidal-ramp protocol (G) to measure the threshold for AP firing at a range of frequencies (data shown as mean ± SEM, n = 7 corpuscles). No difference between the intact (control) and ruptured outer core conditions was detected (two-way repeated measures ANOVA, p=0.1163).

### Direct stimulation of the inner core recapitulates intact corpuscle function

Earlier studies used extracellular recordings to evaluate the adaptation properties of mechanotransduction in the afferent terminal (Bell *et al*., 1994; Loewenstein and Mendelson, 1965; Loewenstein and Skalak, 1966; Mendelson and Loewenstein, 1964). However, extracellular recordings do not fully recapitulate the precise intracellular kinetics of mechanically activated current. To our knowledge, direct evidence of mechano-electric conversion via mechanically gated ion channels in the form of voltage-clamp recordings in the afferent terminal has not been reported for the Pacinian corpuscle in any species. Though the afferent terminal membrane is relatively inaccessible due to being located at the center of the end-organ, we successfully patch-clamped the terminal at the non-myelinated heminode within the corpuscle after breaking through the outer core (Figure 4A). The gap junction-permeable fluorescent dye Lucifer Yellow diffused from the electrode solution along the entire length of the terminal, confirming intracellular access via the patch-clamp electrode. Notably, Lucifer Yellow remained confined within the afferent, suggesting the absence of the dye-permeable gap junctions between the afferent and surrounding cells (Figure 4B). In the afferent terminal, we measured a resting membrane potential of -64.28 ± 1.52 mV, whole-cell capacitance of 11.36 ± 2.04 pF, and input resistance of 238.6 ± 85.96 MΩ (mean ± SEM, n = 5). Mechanical stimulation applied through the outer core, while current-clamping the afferent terminal, elicited action potentials in the terminal, whereas in the voltage-clamp mode it evoked mechanically activated (MA) current during the ON and OFF dynamic phases (Figure 4C, D), in agreement with the rapidly adapting firing recorded in the afferent outside the corpuscle (Figure 3C). These experiments reveal mechanotransduction events at the physiological site of touch detection in the Pacinian corpuscle.

**Figure 4.**
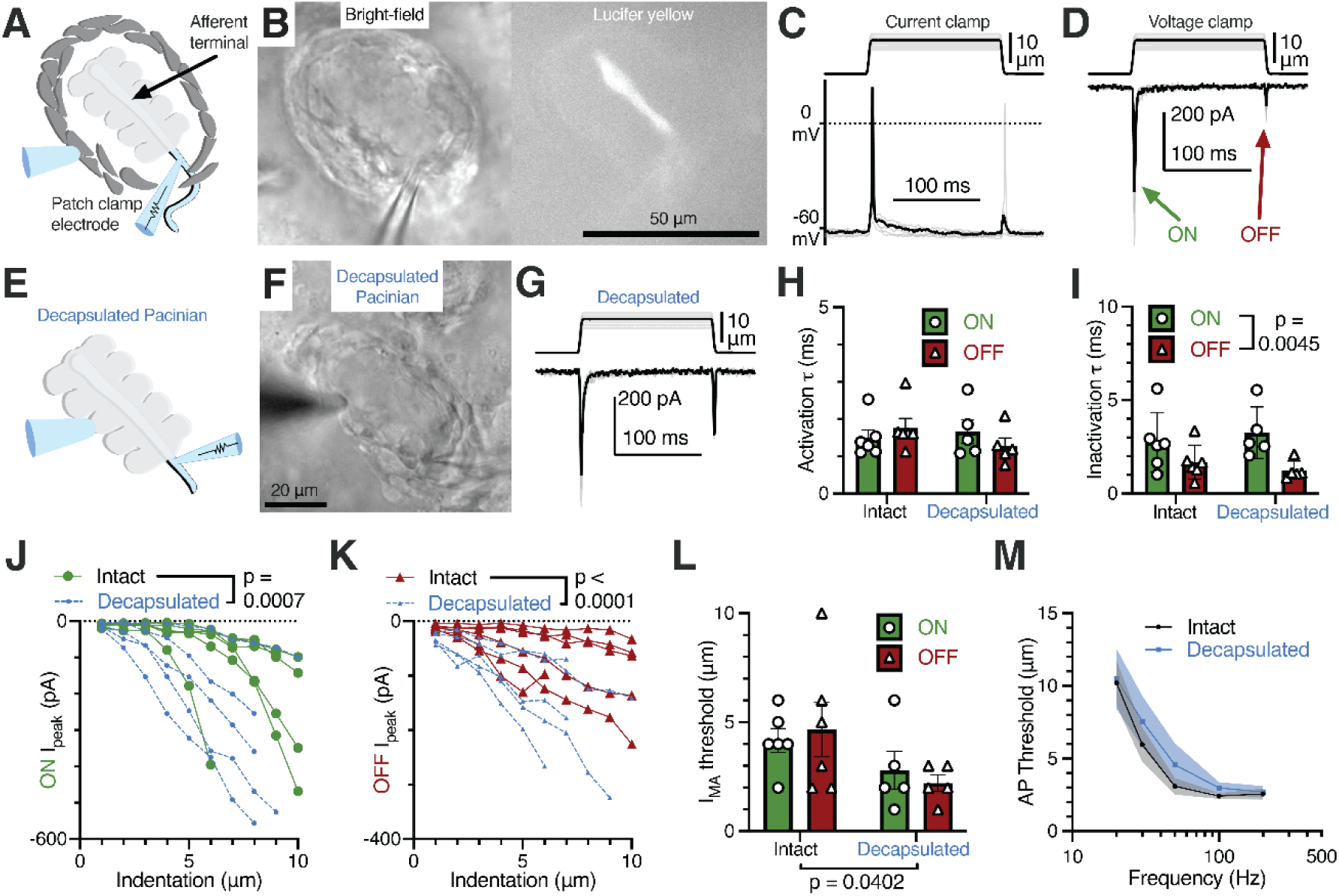
Rapid adaptation and frequency tuning of the Pacinian terminal is independent of the outer core. (A) Illustration of the patch-clamp recording approach of the Pacinian afferent terminal. (B) Bright-field image of the experimental setup under the microscope (left) and lucifer yellow fluorescence in the afferent terminal alone (right). (C) Recordings with the mechanical step stimulus applied with a glass probe (top) and exemplar voltage responses and action potentials (APs) in the terminal in current-clamp mode (bottom). (D) The mechanical stimulus (top) and representative mechanically activated (MA) current responses in the terminal while voltage-clamped at –60 mV (bottom). (E) Illustration of patch-clamp recordings of the terminal of a decapsulated Pacinian corpuscle. (F) Bright-field image of a decapsulated Pacinian. (G) The mechanical stimulus (top) and exemplar MA current responses in a decapsulated Pacinian terminal (bottom). (H, I) Quantification of the kinetics of MA current response activation (H) and inactivation (I) during the ON and OFF phases in intact and decapsulated Pacinian corpuscles. Symbols are values from individual corpuscles. Data shown as mean ± SEM. Statistics: two-way ANOVA. No difference was detected between intact and decapsulated activation (p=0.5409) nor inactivation (p=0.9418) (J, K) The peak current recorded in the ON (J) and OFF (K) phases in relation to the indentation depth of the probe. Each line represents one cell. Statistics: two-way ANOVA. (L) Comparison of the current response threshold of the ON and OFF phases between terminals of intact and decapsulated corpuscles. Symbols are values from individual corpuscles. Data shown as mean ± SEM. Statistics: two-way ANOVA. (M) Population tuning curve of intact and decapsulated Pacinians measured via single-fiber recording. Data shown as mean ± SEM from 6 corpuscles. No effect of decapsulation was detected (two-way repeated measures ANOVA, p=0.3842).

To directly test if the outer core influences afferent terminal mechanotransduction, we sought to compare functional properties of MA current in the terminal in response to stimulation applied either to the outer core or directly to the inner core. To this end, we mechanically decapsulated the corpuscle by removing the outer core such that the inner core became accessible to direct mechanical stimulation (Figure 4E, F). Strikingly, and in agreement with our experiments above, we found that MA currents in intact and decapsulated corpuscles were largely indistinguishable. In both conditions MA currents occurred only during the dynamic phases of the stimulus (Figure 4G) and retained equally fast kinetics of activation and inactivation (Figure 4H, I). However, after decapsulation we observed an increase in MA current amplitude (Figure 4J, K) and a concurrent decrease in the threshold of activation for both ON and OFF responses (Figure 4L). Thus, our results support the idea that while the outer core may present a physical layer that attenuates the threshold of mechanical stimulus detection, it does not influence the timing of activation and inactivation of MA current in the afferent terminal.

Earlier reports showed that the removal of the outer core profoundly increases the time of receptor potential decay recorded in a Pacinian afferent (Bell *et al*., 1994; Loewenstein and Mendelson, 1965; Loewenstein and Skalak, 1966; Mendelson and Loewenstein, 1964). These studies suggested that mechanically gated ion channels in the terminal have slow kinetics of inactivation, and that they only appear fast due to the presence of the outer core, which acts as a mechanical filter that limits static stimuli from reaching the terminal. In contrast, our data show that MA current kinetics remains fast regardless of the presence of the outer core. We explored a possible cause for this apparent contradiction. We noticed that decapsulation leads to a visible deterioration of the afferent terminal, and 10 minutes after decapsulation the terminal begins to show MA current with slow inactivation kinetics (Figure S7A-D). Furthermore, slower inactivation rates were correlated with larger currents required to hold the terminal at -60 mV (Figure S7E), demonstrating that terminals in declining health display prolonged MA current decay. Thus, our data agree with the idea that the outer core provides an optimal environment for the inner core function, but its influence is independent of its proposed role as a mechanical filter.

The fast kinetics of MA current inactivation in the Pacinian afferent terminal prompted us to perform a comparison with duck Meissner (Grandry) corpuscles. The tuning curve of Meissner corpuscles has a peak sensitivity at a lower frequency than that of Pacinians, in agreement with the functional specialization of these end-organs in mammals and birds as detectors of, respectively, low and high frequency vibration (Figure S8A) (Ziolkowski *et al*., 2022). Consistently, the inactivation rate of MA current in the Pacinian terminal is significantly faster than in the Meissner terminal (Figure S8B, C) (Ziolkowski *et al*., 2023), supporting the notion that faster inactivation is more conducive to detecting high frequency vibration.

Next, we tested whether the outer core is required for high-pass frequency filtering by recording the discharge of the Pacinian afferent in the same corpuscle before and after decapsulation. We found that direct stimulation of the inner core failed to affect frequency tuning (Figure 4M). Together, our data show that while the outer core likely serves a protective role by providing the optimal environment for the inner core, it is not necessary for rapid adaptation of the afferent discharge, fast inactivation of MA current, or frequency tuning. Instead, these key functions of Pacinian corpuscles originate from the inner core.

### Inner core lamellar Schwann cells form a syncytium coupled via gap junctions

Given the localization of primary corpuscle function to the terminal and inner core, we next sought to investigate the electrophysiological properties of LSCs. To our knowledge electrophysiological recordings from these cells have not been reported for any species. As we report here, inner core lamellae of duck Pacinian corpuscles are connected by gap junctions (Figure S6), suggesting that LSCs may influence the electrical properties of adjacent LSCs, but whether gap junctions functionally couple these cells is unknown. We found that Lucifer Yellow injected into a single lamellar cell via the recording electrode diffused among all LSCs, however it did not spread into the afferent or other cells outside the inner core (Figure 5A, B), which demonstrates free passage of small molecules between LSCs. Subsequent patch-clamp measurements from LSCs revealed a resting membrane potential of -73.01 ± 1.664 mV (mean ± SEM, n = 20). In agreement with the presence of extensive lamellae and intracellular connections formed between LSCs, these cells exhibit unusually high values of whole cell capacitance (425.0 ± 33.8 pF, n = 16) as measured via a negative voltage step (Figure 5C, D). Accordingly, RNA sequencing of inner cores individually extracted from Pacinian corpuscles showed robust expression of gap junction proteins, including various types of connexins (Figure S9 and Supplementary Data S1). To further test gap junction coupling, we measured membrane capacitance and input resistance before and after adding the pan-gap junction blocker carbenoxolone (CBX, Figure 5C-E). The addition of CBX reduced whole cell capacitance 12-fold to 36.7 ± 8.9 pF (Figure 5D) and doubled the input resistance from 568 ± 86 MΩ to 1,116 ± 177 MΩ (Figure 5E), supporting the idea of functional gap junction coupling between LSCs.

**Figure 5.**
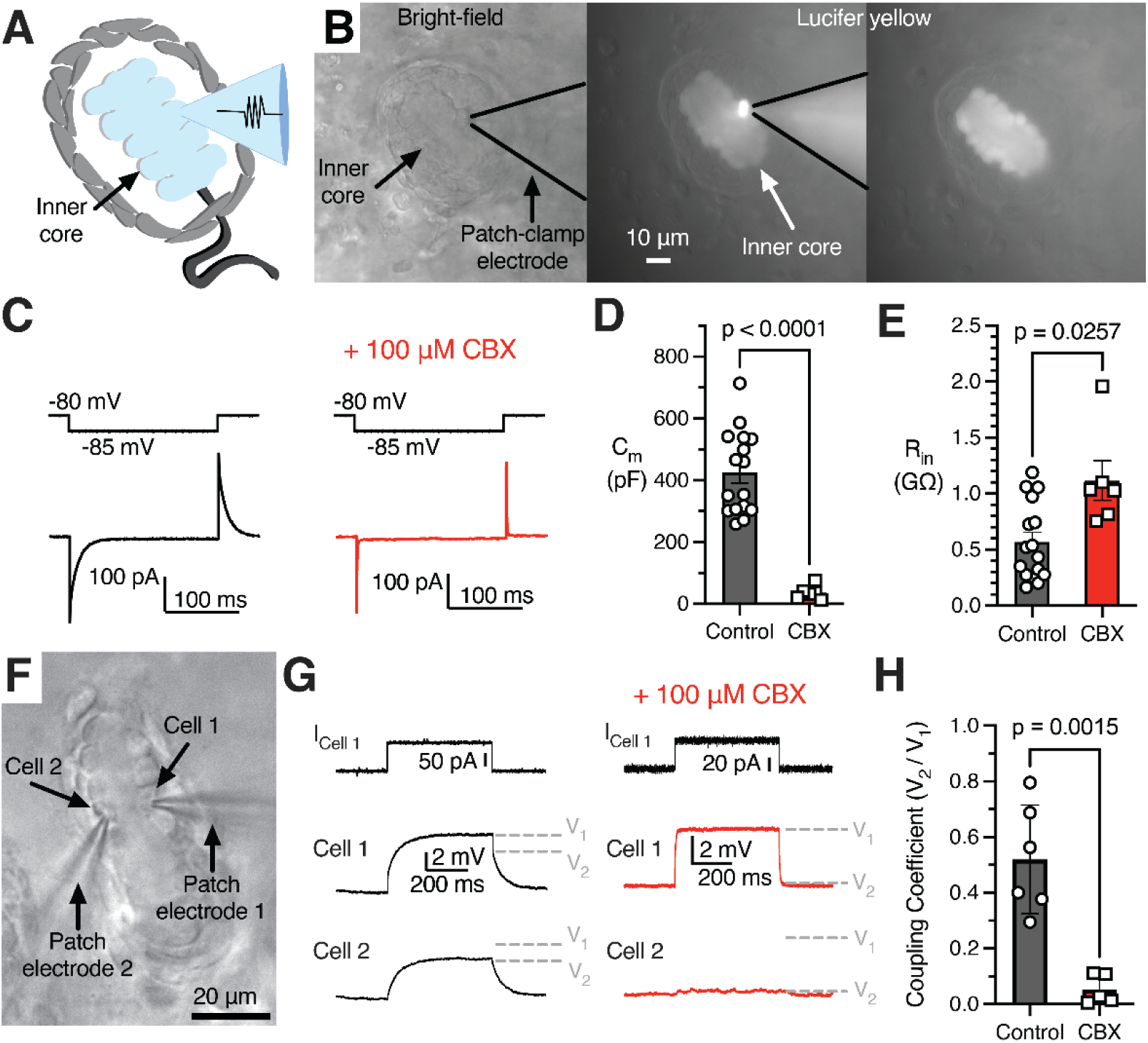
Lamellar Schwann cells form a gap junction-coupled syncytium. (A) Illustration of interconnected, patch-clamped inner core LSCs. (B) Bright-field image of a patched LSC (left), fluorescence of Lucifer Yellow in the patched inner core (middle), and fluorescence in the inner core after the patch electrode is removed (right). (C) A small voltage step applied to the patched LSC (top) and the example current responses (bottom) with normal intracellular solution or including the gap junction blocker CBX (right). (D, E) Membrane capacitance (D) and input resistance (E) with and without CBX. Symbols are recordings from individual corpuscles. Data shown as mean ± SEM. Statistics: Welch’s t-test. (E) Input resistance with and without CBX. (F) Bright-field image of simultaneous dual LSC patch clamp recordings. (G) Simultaneous recordings from adjacent LSCs (Cell1 and Cell 2). A current injection stimulus applied to Cell 1 in the recording setup in F (top), an example voltage response in Cell 1 (middle), and an example voltage response in the connected Cell 2 (bottom), with normal intracellular solution or including CBX. V_1_ and V_2_ denote voltage levels induced in, respectively, Cell 1 and Cell 2. (H) The coupling coefficient between two cells in control conditions *vs.* with CBX. Symbols are recordings from individual corpuscles. Data shown as mean ± SEM. Statistics: Welch’s t-test.

Next, we quantified electrical coupling of two LSCs near resting membrane potential by performing dual patch-clamp recordings (Figure 5F). A current injection stimulus applied to the first LSC elicited a voltage response in the same cell, and a smaller voltage response in the second cell, which was abolished by adding CBX to block gap junctions (Figure 5G). The coupling coefficient between LSCs, quantified as the ratio of the voltage response between the two cells, was found to be 0.52 ± 0.08. This value significantly reduced in the presence of CBX to 0.05 ± 0.02 (Figure 5H). However, the coupling coefficient progressively decreased upon larger current injections, demonstrating that electrical coupling between adjacent LSCs is voltage-dependent and is efficient only for small-scale depolarizations (Figure S10). These results demonstrate that gap junctions connect the cytoplasm of adjacent LSCs, permitting a free flow of small molecules between the cells and forming an electrically coupled syncytium. Because gap junctions were also reported earlier between inner core lamellae of Pacinians from dogs and cats (Ide and Hayashi, 1987; Rico et al., 1996), and in the accompanying study in mice (Chen *et al*.) our findings suggest that the electrical coupling between LSCs could be an evolutionary conserved feature of Pacinian corpuscles.

### Inner core lamellar Schwann cells are mechanosensitive

Our RNA sequencing of Pacinian inner cores revealed expression of various types of known mechanically gated ion channels and their modifiers (Kefauver et al., 2020; Syeda, 2021; Zhou et al., 2023), and voltage-gated ion channels (Figure S9B, C). We hypothesized that LSCs could be excitable mechanosensors that actively participate in touch detection. We tested this by applying mechanical stimuli to patch-clamped LSCs (Figure 6A) and found that they indeed responded with robust MA current (Figure 6B), which increased in magnitude with larger indentation depths (Figure 6C). MA current displayed virtually no inactivation during the 150 ms long stimulation and exhibited a noticeable persistent component (26.51% ± 3.57% of the peak current) that remained after the indentation probe was retracted. Mechanical stimulation to the same depths at a range of voltages revealed a linear voltage-dependence of MA current with a reversal potential of 16.40 mV (95% confidence intervals of 6.25 to 28.06 mV, Figure 6D, E), indicative of conductance with poor ionic selectivity.

**Figure 6.**
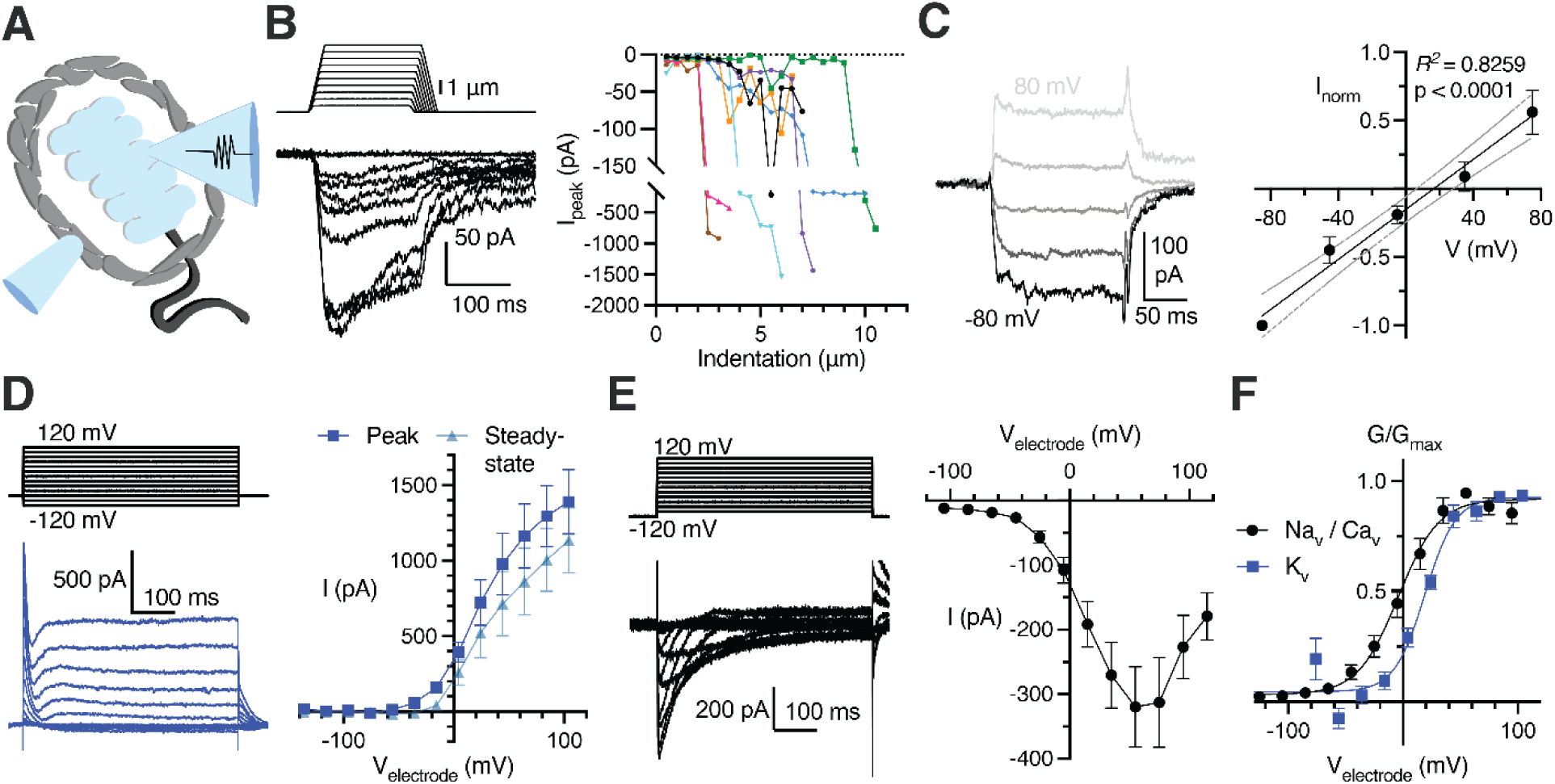
Lamellar Schwann cells are mechanosensitive and express voltage-gated ion channels. (A) Illustration of a patch-clamped LSC with an indentation probe for mechanical stimulation (B) Mechanical step stimuli (top left) and example voltage-clamp recordings (bottom left) from LSCs held at -80 mV, displaying inward mechanically activated (MA) current, and the summary of peak MA current *vs.* indentation depth in LSCs (right). Lines connect data from individual LSCs. (C) Example MA current responses in a LSC voltage-clamped at -80, -40, 0, 40, and 80 mV and the current-voltage relationship (I-V curve) of MA current (mean ± SEM, n = 6 LSCs). (D) The voltage step stimulus and an exemplar voltage-gated current response from an LSC with potassium-based internal solution, displaying heterogenous outward voltage-gated current (K_v_ current) (left) and the I-V curve of the K_v_ current measured at peak or steady-state phase of the current (right). Data shown as mean ± SEM from 11 recordings. (E) The voltage step stimulus and an exemplar voltage-gated current response from an LSC with cesium-based internal solution, displaying inward, inactivating voltage-gated current (Na_v_ or Ca_v_ current) (left) and the I-V curve of the inward voltage-gated current (right). Data shown as mean ± SEM from 10 recordings. (F) Conductance-voltage relationship of the Na_V_/Ca_V_ and K_V_ current, fitted with the Boltzmann equation. Data shown as mean ± SEM from 10 Na_v_/Ca_v_ and 11 K_v_ recordings.

Next, we tested whether LSCs express voltage-gated ion channels. We applied a voltage-step protocol with potassium-based internal solution and detected voltage-activated outward currents (Figure 6F, G, J), demonstrating the presence of voltage-gated potassium (K_V_) channels. After blocking K_V_s with cesium-based internal solution, we detected voltage-gated inward currents (Figure 6H-J), revealing the presence of voltage-activated sodium (Na_V_) and/or calcium (Ca_V_) channels in LSCs. These data are consistent with the expression of various types of K_V_, Na_V_ and Ca_V_ transcripts in our RNA sequencing of Pacinian inner cores (Figure S9C). The voltage dependence of activation of Na_V_/Ca_V_ conductances appear stretched towards positive potentials in comparison with known Na_V_ and Ca_V_ channels *in vitro* and in native cells (Catterall, 2023), suggesting that many of these channels are expressed in the long lamellar processes of LSCs, and the recorded right-shifted voltage dependance likely reflects voltage drop produced by the high electrical resistance of these structures. Together, these data show that LSCs are mechanosensitive and express depolarizing and hyperpolarizing voltage-gated ion channels.

### Activation of inner core lamellar Schwann cells increases afferent sensitivity to touch

We were ultimately led to ask whether LSCs affect the function of the mechanoreceptor terminal, which thus far has been thought to be the sole site of touch detection in Pacinian corpuscles. We first performed single-fiber recording of the Pacinian afferent with simultaneous patch-clamp stimulation of an LSC, but we were not able to induce any AP firing in the afferent upon activation of LSCs by current injection or depolarization (Figure S11). Because single-fiber recording may fail to detect sub-threshold intracellular responses in the afferent, we performed voltage-clamp recordings of the mechanoreceptor terminal paired with simultaneous stimulation of an LSC (Figure 7A). Using this setup, we detected a depolarizing inward current in the afferent terminal in response to activation of an LSC with current injection (Figure 7B). In contrast, activation of an LSC failed to induce current in an adjacent OCLC from the outer core, demonstrating that the functional coupling was specifically between LSCs and the mechanoreceptor terminal (Figure 7C). Consistently, in current-clamp mode, the afferent terminal responded to LSC activation by depolarization, which increased with larger current injection into the LSC (Figure 7D, E).

**Figure 7.**
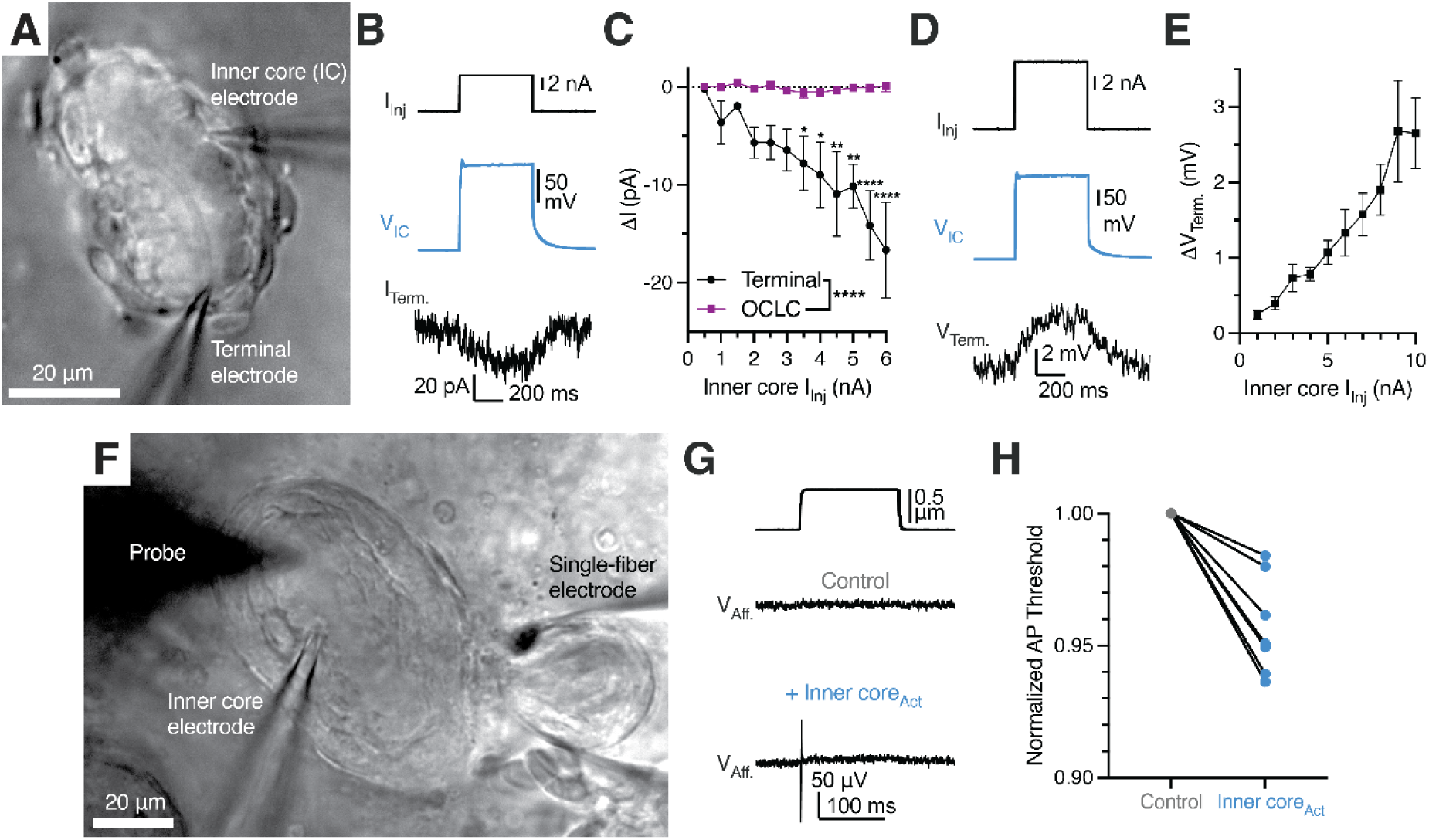
Activation of lamellar Schwann cells reduces the mechanosensitivity threshold of the Pacinian corpuscle. (A) Bright-field image of simultaneous paired patch clamp recordings from one LSC and an associated Pacinian afferent of the same corpuscle. (B) Exemplar traces showing current injection stimulus applied to a LSC (top), voltage response of the LSC (middle) and current response of the afferent terminal voltage-clamped at -60 mV (bottom). (C) Quantification of current responses in the afferent terminal and an OCLC upon current injection into an LSC. Data shown as mean ± SEM from 4 afferent terminal and 4 OCLCs recordings. Statistics: two-way ANOVA with Holm-Šidák post-hoc tests (*p<0.05, **p<0.01, ****p<0.0001) (D) Exemplar traces showing a current injection stimulus applied to a LSC (top), voltage response of the LSC (middle) and voltage response of the current-clamped afferent terminal (bottom). (E) Quantification of voltage response in the afferent terminal upon current injection into an LSC. Data shown as mean ± SEM from 7 recordings. (F) Bright-field image of simultaneous paired patch clamp recordings from one LSC and single-fiber recording of an associated Pacinian afferent of the same corpuscle while applied mechanical stimuli with the marked probe to measure AP threshold. (G) Mechanical stimulus (top) and single-fiber recordings of the Pacinian afferent during the absence (middle) or presence (bottom) of LSC-inner core activation via 6 nA current injection (H) Quantification of the effect of LSC activation by depolarizing current injection (6 nA) on the threshold of mechanical activation in Pacinian afferent evoked by to a square indentation step. Lines connect data from individual paired recordings. Mechanical threshold during inner core activation was lower than the normalized control threshold (p = 0.001, one sample t-test).

Having established that LSCs can influence the excitatory status of the afferent, we hypothesized that LSC-induced depolarization of the afferent terminal should lower the threshold of mechanical stimulation required for the mechanoreceptor to generate an action potential. Indeed, we found that activation of an LSC by current injection triggered AP firing in the afferent in response to a sub-threshold mechanical stimulus (Figure 7F), and lead to an overall reduction of the threshold required for activation of the afferent by mechanical force (Figure 7G). In agreement with our data, the accompanying study shows that optogenetic inhibition of LSCs in mouse Pacinian corpuscles increases the threshold of mechanical activation (Chen *et al*.). Together, our works establish an evolutionarily conserved role of LSCs as active mechanosensory elements within Pacinian corpuscles that potentiate sensitivity of the afferent terminal to mechanical touch.

## Discussion

Our results demonstrate that, contrary to the accepted view, the outer core is dispensable for rapid adaptation and frequency tuning – the main functional properties of Pacinian corpuscles. We observe that when the integrity of the outer core is compromised, or when mechanical stimulation is delivered to the inner core directly, bypassing the outer core, the afferent nevertheless displays rapid adaptation and high-pass frequency filtering. This response, which is characteristic of mature mammalian (including human) and avian Pacinian corpuscles, is indistinguishable from that obtained by stimulation of an intact end-organ. Earlier observations using Pacinian corpuscles from cat mesentery documented that physical removal of most outer core layers converts the timing of mechanically evoked receptor potential decay from fast to slow (Loewenstein and Mendelson, 1965; Loewenstein and Rathkamp, 1958a; b; Mendelson and Loewenstein, 1964). Other studies, however, noted that this only happens upon intense compression, whereas light forces do not affect the process (Hunt and Takeuchi, 1962; Nishi and Sato, 1968; Ozeki and Sato, 1965). These observations suggested a model in which the outer core acts as a multi-layered mechanical cushion that prevents static stimuli from reaching the core (Loewenstein and Skalak, 1966). This model was further extended to suggest that the same mechanism could be responsible for high-pass frequency tuning (Bell *et al*., 1994; Quindlen-Hotek *et al*., 2020; Quindlen *et al*., 2016), even though, to our knowledge, this idea has not been tested until now via a comparison of intact *versus* decapsulated corpuscles. While our experiments demonstrate that direct mechanical stimulation of isolated inner cores produces the same functional outcome as stimulation of the intact structures, they also show that over time, as the afferent terminal deteriorates, the timing of MA current decay becomes noticeably slow. Our observations thus agree with the idea that the outer core provides a protective environment for the inner core components (Gray and Sato, 1955; Ilyinsky *et al*., 1976), but rapid adaptation and frequency filtering are mediated by the inner core independently of the outer core.

Despite differences in the overall size, avian and mammalian Pacinian corpuscles share the same overall topology, including the presence of an outer core, inner core and afferent terminal with protrusions. The 3D structures of inner cores shown here for the avian Pacinian corpuscle and in the accompanying study for its mouse counterpart (Chen *et al*.) permit a detailed comparison of both structures. In both cases, the inner core is composed of two columns of LSCs which extend long interdigitating lamellae, encompassing the entire length of the afferent terminal. In both cases, the afferent terminal contains thin protrusions which penetrate through inner core lamellae and extend to the periphery. Although the number of protrusions in the avian Pacinian is significantly smaller than in mouse corpuscles, which could be either species-specific or reflect a developmental stage, they share a similar overall appearance and similarly originate from opposite sides of the elliptical terminal. Because protrusions are present in Pacinians from different species (Bolanowski *et al*., 1994; Handler *et al*., 2023; Zelena *et al*., 1997) they are a general feature of Pacinian corpuscles. While the exact function of the protrusions remains to be determined, they were proposed to be the key sites of mechanotransduction in the terminal (Bolanowski *et al*., 1994), and in mice were shown to express the Piezo2 ion channel (Handler *et al*., 2023).

Because of the structure and location, the individual components of Pacinian corpuscles are nearly impervious to direct electrophysiological investigations. Here, we recorded via patch-clamp the afferent terminal within the corpuscle to reveal biophysical properties of the underlying mechanically gated ion channels. We found that both ON and OFF MA currents exponentially decay with remarkably fast kinetics. Our measured inactivation constant (τ_inact_ = 1-3 ms) is smaller than those previously recorded in mechanoreceptor afferents in worms (Das et al., 2024; Eastwood et al., 2015; Katta et al., 2019; O’Hagan et al., 2005), in Merkel afferents from mouse whiskers (Yamada et al., 2024), and in Meissner corpuscle afferents in ducks (Ziolkowski *et al*., 2023). In mammals, afferents innervating Merkel cells, Meissner and Pacinian corpuscles express Piezo2 (Garcia-Mesa et al., 2024; Garcia-Mesa et al., 2022; Handler *et al*., 2023; Ranade et al., 2014), and it is likely to be the major ion channel mediating MA current in avian afferents (Schneider et al., 2019; Schneider et al., 2014). Although the MA current in Pacinian afferents inactivates much faster than mouse or duck Piezo2 *in vitro* (τ_inact_ ≈ 8-10ms) (Coste et al., 2010; Schneider et al., 2017), Piezo2 inactivation is a variable parameter influenced by cell-specific factors (Anderson et al., 2018; Del Rosario et al., 2022; Dubin et al., 2012; Ma et al., 2023; Romero et al., 2023; Romero et al., 2020; Schaefer et al., 2023; Zhang et al., 2024; Zheng et al., 2019a; Zheng et al., 2019b; Zhou *et al*., 2023).

The remarkable architecture of the Pacinian inner core presented here and in the accompanying study (Chen *et al*.) reveals interdigitating crescent-shaped lamellae formed by LSCs that envelop the afferent terminal. This raises the question of whether and how the lamellae formed by LSCs contribute to corpuscle function. One possibility is that the inner core lamellae perform the ‘mechanical filter’ role previously assigned to the OCLC layers in the outer core. Indeed, the sensory Schwann cells in mammalian Meissner corpuscles, which exhibit frequency tuning and rapid adaptation, also sprout lamellae around the afferent terminal, but are devoid of an outer core (Handler *et al*., 2023). Avian Meissner corpuscles also contain sensory Schwann cells (Nikolaev *et al*., 2020), which are transcriptionally similar to Pacinian LSCs (Figure S12). Like their mammalian counterparts, avian Meissner corpuscles show rapid adaptation and frequency tuning, but their sensory Schwann cells do not form lamellae around the afferent (Nikolaev *et al*., 2023). These observations suggest that a lamellar structure around the afferent is, in principle, not essential for rapid adaptation and frequency tuning.

We show that mechanically gated ion channels in the Pacinian afferent terminal open during the dynamic phases of mechanical stimulus, causing AP firing, but then quickly close and remain inactivated during the static phase. We made similar observations in the afferent terminal of avian Meissner corpuscles (Ziolkowski *et al*., 2023), indicating that rapid adaptation is likely an inherent consequence of fast channel inactivation. This conclusion may extend to frequency filtering of corpuscles, as a recent study showed that the efficiency of Piezo2 activation increases with indentation velocity, suggesting that the channel is more effectively engaged by high frequency stimulation (Zeitzschel and Lechner, 2024). Interestingly, the significantly faster inactivation of MA current in the Pacinian terminal compared to the Meissner terminal shown here could additionally explain why these structures are tuned to higher and lower frequencies, respectively. Our observations thus support the idea that rapid adaptation and frequency filtering could stem from biophysical properties of mechanotransducing ion channels in the afferent terminal. However, the accompanying study demonstrates that LSCs play a critical role in shaping the frequency filtering of mouse Pacinian corpuscles (Chen *et al*.). Together, these data suggest a general model where the interplay between sensory Schwann cells of Meissner corpuscles and in the Pacinian inner core influence inactivation rates of the mechanically gated ion channels in the afferent terminal to control precise adaptation rates and frequency filtering.

Here, we showed that LSCs express slowly inactivating mechanically gated ionic conductance. Although the molecular identity of slowly inactivating channels in these cells is unknown, our finding establishes LSCs as mechanosensors. Moreover, because activation of an LSC decreases the threshold of mechanical activation of the Pacinian afferent, our data establish LSCs as physiologically relevant touch sensors which facilitate mechanosensitivity of Pacinian corpuscles. The accompanying study shows that this is also true for mouse Pacinians (Chen *et al*.), demonstrating that the multicellular mechanism of touch detection is evolutionarily conserved in Pacinian corpuscles from different species.

How Pacinian LSCs facilitate mechanosensitivity of the afferent terminal remains an open question. Our observations rule out direct electrical coupling between LSCs and the terminal via gap junctions. The absence of clearly identifiable vesicles in LSCs or synapse-like structures between LSCs and the terminal also argues against, though does not rule out, synaptic-like mechanisms as reported for Merkel cell-neurite complexes (Chang et al., 2016; Hoffman et al., 2018; Yamada *et al*., 2024). It is possible that other mechanisms, such as ephaptic cross-talk between adjacent membranes, or ion channel-based communication, such as those mediating interaction between peripheral glia and mechanoreceptors in worms (Fernandez-Abascal et al., 2021; Graziano et al., 2024) and between keratinocytes and mechanoreceptors in mice (Moehring et al., 2018) are at play. Prominent tethers connecting LSC lamellae with afferent membrane suggest yet another possibility of physical coupling between the lamellae and mechanically gated channels in the terminal (Das *et al*., 2024; Hu et al., 2010; Li and Ginty, 2014; Nikolaev *et al*., 2023; Schwaller et al., 2021). It remains to be determined if any of these mechanisms partake in Pacinian corpuscle function, and whether they are present in other mechanoreceptive end-organs in vertebrates, which contain sensory Schwann cells (Abdo et al., 2019; Handler *et al*., 2023; Nikolaev *et al*., 2023; Ojeda-Alonso et al., 2024; Qi et al., 2024).

## Acknowledgements

We thank members of the Bagriantsev, Gracheva and Daniel Huber’s laboratories for comments and critique throughout the study; Morven Graham, Xinran Liu and Yale School of Medicine Electron Microscopy Core for transmission electron microscopic imaging for electron tomography; Wei-Ping Li for the support of EM sample preparation for eFIB-SEM imaging, Benjamin Bae for help with eFIB-SEM data processing, Jazune Madas, Argaja Shende, and Chris Zugates for advice on eFIB-SEM data segmentation, FIB-SEM Collaboration Core at Yale School of Medicine for enhanced FIB-SEM pipeline support.

## Author contributions

L.H.Z, Y.A.N., A.C., M.O., V.V.F., C.S.X., S.P. conducted experiments. L.H.Z, Y.A.N., A.C., M.O., V.V.F., D.M.-A., S.A.A., S.S.B. analyzed data. C.S.X supervised eFIB-SEM experiments. E.O.G., S.N.B. conceived and supervised the project. L.H.Z., Y.A.N., E.O.G., S.N.B. wrote the manuscript with input from all authors.

## STAR Methods

### Animals

Experiments with Mallard duck embryos (*Anas platyrhynchos domesticus*) were approved by and performed in accordance with guidelines of the Institutional Animal Case and Use Committee of Yale University (protocol 11526). Animals used in experiments were at development stages embryonic day 25 (E25) to E27, between 1-3 days before hatching; sex was not determined.

### *Ex vivo* bill-skin preparation

Dissection of bill-skin was performed as described previously (Nikolaev *et al*., 2023; Ziolkowski *et al*., 2023). First, the glabrous skin of the bill was shaved off from the embryo and put into ice-cold L-15 media, where it was trimmed to fit into a recording chamber. For experiments involving single-fiber or patch-clamp recording of the afferent, the bill-skin was inverted (with the dermis on top and epidermis on the bottom) in the recording chamber in Krebs solution containing (in mM) 117 NaCl, 3.5 KCl, 2.5 CaCl_2_, 1.2 MgCl_2_, 1.2 NaH_2_PO_4_, 25 NaHCO_3_, and 11 glucose, saturated with 95% O_2_ and 5% CO_2_ (pH 7.3-7.4), at room temperature (22-23°C). The bill-skin was treated with 2 mg/mL collagenase P (Roche) in Krebs solution for 5 minutes, then washed with fresh Krebs solution. Bill-skin preparations used for solely lamellar cell patch-clamp recording were placed into Ringer solution containing (in mM) 140 NaCl, 5 KCl, 10 4-(2-hydroxyethyl)piperazine-1-ethane-sulfonic acid (HEPES), 2.5 CaCl_2_, 1 MgCl_2_, and 10 glucose at room temperature. The epidermis was carefully removed from the dermis, which was treated with 2 mg/mL collagenase P in Ringer solution for 5 minutes, then washed with fresh Ringer. Corpuscles in the dermis were visualized on an Olympus BX51WI upright microscope with an ORCA-Flash 4.0 LT camera (Hamamatsu).

### Electrophysiology

#### Single-fiber recordings from individual Pacinian afferents

Recordings from single afferent fibers of avian Pacinian (Herbst) corpuscles were acquired at room temperature in Krebs solution using a MultiClamp 700B amplifier and Digidata 1550A digitizer (Molecular Devices). Single-fiber recording pipettes were created from borosilicate glass capillaries with outer diameter 1.5 mm, inner diameter 1.17 mm, wall thickness 0.17 mm, without filament (Warner Instruments model GC150T-7.5). Pipettes were pulled using a P-1000 micropipette puller (Sutter Instruments) to create tip diameters of 5 to 30 μm, then filled with Krebs solution. Pipettes were placed on a CV-7B headstage connected to a High-Speed Pressure Clamp (ALA Scientific Instruments). Single corpuscles and connected afferents within the same field of view were identified under a 40X objective lens. The recording pipette was placed next to the afferent, and negative pressure was applied until a large section (∼5 μm) of the afferent was sucked into the pipette. The extracellular afferent voltage was recording in current-clamp mode, sampled at 20 kHz and low-pass filtered at 1 kHz in Clampex 10.7 (Molecular Devices). A suprathreshold mechanical step stimulus was applied to the connected corpuscle to confirm the presence of mechanically induced action potentials in the afferent fiber. Fresh Krebs solution was regularly perfused onto the preparation between recordings.

Mechanical stimuli were applied to a single corpuscle using a blunt glass probe (5 to 10 μm tip diameter) mounted on a piezoelectric-driven actuator (Physik Instrumente Gmbh). A mechanical step stimulus was applied to corpuscles with variable displacements in increments of 1 μm. The duration of the static and dynamic phases of the step stimulus were constant at 150 ms and 3 ms, respectively. Vibratory stimuli were applied using a sinusoidal-ramp waveform, increasing 0.25 μm per cycle, at frequencies of 20, 30, 50, 100, 200, and 400 Hertz. AP threshold was defined as the smallest probe displacement which elicited an action potential. To block APs, 1 μM tetrodotoxin citrate (Tocris) was added to the bath. For experiments with ruptured corpuscles, a puncture was created in the outer core using a high-pressure stream of Krebs solution from a patch pipette.

#### Patch-clamp electrophysiology

Whole-cell recordings of afferent terminals and lamellar Schwann cells were performed at room temperature, using the same amplifier and digitizer used for single-fiber recording. Borosilicate pipettes with filament, outer diameter 1.5 mm, inner diameter 0.86 mm, wall thickness 0.32 mm, and tip resistances of 2-7 MΩ were used to acquire voltage-clamp and current-clamp recordings. Unless otherwise indicated, pipettes were filled with potassium-based internal solution (K-internal) containing (in mM) 135 K-gluconate, 5 KCl, 0.5 CaCl_2_, 2 MgCl_2_, 5 EGTA, 5 HEPES, 5 Na_2_ATP, and 0.5 Na_2_GTP (pH 7.3 with KOH) and placed on a CV-7B headstage connected to a High-Speed Pressure Clamp. In certain experiments, 1 mM Lucifer yellow (Sigma-Aldrich) was included in the internal solution and fluorescently excited with a U-HGLGPS illumination source (Olympus) and Lucifer yellow filter cube to visualize the patched cell. In almost all experiments, Krebs was used as the external bath solution. For patch-clamp recordings from LSCs, Ringer solution was used as the external solution. Data from intracellular recording was sampled at 20 kHz and low pass filtered at 2 kHz in Clampex 10.7. Paired recordings from a second cell were acquired with a CV-7B headstage connected to the other channel of the same amplifier and digitizer, using the same single-fiber or patch-clamp techniques described here.

To access the membrane of the afferent terminal and LSCs, large positive pressure (>100 mmHg) was applied to the recording pipette, which was used to pierce the outer core of the corpuscle and remove obstructions blocking the desired cell. In decapsulation experiments, the inner core was separated entirely from the outer core using this method. Occasionally, multiple patch pipettes were used to blow debris away from the target cell membrane before sealing and break-in. For afferent terminal voltage-clamp recordings, the cells were clamped at -60 mV and the same mechanical stimuli applied during single-fiber recording were used. Only data from healthy terminals (holding currents above -75 pA during voltage-clamp at -60 mV) were used in analysis, except to explore the relationship between holding current and MA current inactivation rate, in which unhealthy terminals (holding current below -75 pA) were included. The inactivation rate (τ) of the MA current was calculated as described previously (Nikolaev *et al*., 2020) by fitting a single exponential function (I = I_0_*exp^(− t/τ), where I_0_ is the baseline-subtracted peak current amplitude, t is the time from the peak current, and τ is the inactivation constant) to the decaying portion of the responses in the ON and OFF phases. The MA current threshold was defined as smallest probe displacement that elicited a response in which the amplitude exceeded 20 pA from baseline.

Electrical properties of LSCs were recorded within 30 seconds of establishing whole-cell mode. To block gap junctions in certain experiments, 100 μM carbenoxolone (CBX) was included in the internal pipette solution. For paired LSCs recording, the coupling coefficient was measured as follows: In current-clamp, a 40-100 pA current step was injected into the first cell to elicit a small depolarization (V_1_). The resultant depolarization in the connected second cell (V_2_) was measured, and the ratio V_2_/ V_1_ was calculated. During voltage-clamp experiments, LSCs were clamped at -80 mV unless otherwise indicated. Mechanical ramp-and-hold stimuli were applied to LSCs with increasing static displacement increments of 0.5 μm held for 150 ms, and constant ramp velocities of 1,000 μm/s. MA current was recorded with cesium-based intracellular solution (Cs-internal) containing (in mM) 133 CsCl, 10 HEPES, 5 EGTA, 1 CaCl_2_, 1 MgCl_2_, 4 MgATP, and 0.4 Na_2_GTP (pH = 7.3 with CsOH). Voltage-gated potassium currents were recorded with K-internal including 100 μM CBX to isolate single LSCs. by applying 500 ms depolarizing voltage steps in 20 mV increments (-120 to 120 mV) from -80 mV. Inward Na^+^/Ca^2+^ voltage-gated currents were recorded using Cs-internal to block potassium current along with 100 μM CBX to isolate single LSCs. In this case, 500 ms depolarizing voltage steps in 20 mV increments (-100 to 120 mV) were applied after hyperpolarizing the cell to -120 mV for 2 s to remove channel inactivation. Voltage-activated conductance was calculated using the equation G = I / (V_m_ – E_rev_), where G is the conductance, I is the peak current, V_m_ is the membrane potential and E_rev_ is the reversal potential. The conductance data were fit with the modified Boltzmann equation, G = G_min_ + (G_max_ – G_min_) / (1 + exp^([V_1/2_ – V_m_]/k)), where G_min_ and G_max_ are minimal and maximal conductance, respectively, V_m_ is the voltage, V_1/2_ is the voltage at which the channels reached 50% of their maximal conductance, and k is the slope of the curve. LSC voltage-clamp experiments were corrected offline for liquid junction potential calculated in Clampex 10.7.

Paired recordings were acquired in Krebs solution, with K-internal in patch-clamped cells. Because of the low input resistance of LSCs with open gap junctions, large current injections (1-10 nA) were used to generate voltage responses in the inner core capable of depolarizing the afferent in paired recordings. Mechanoreceptor AP threshold in paired single-fiber/LSC patch-clamp recording was measured via 0.010 μm increments from 0.25 μm range set below and including the baseline threshold determined before the experimental protocol. Two identical mechanical step stimuli of equal displacement were applied, first with the LSC at rest and then with the LSC depolarized by 6 nA for 500 ms starting 150 ms before the second mechanical stimulus. After incrementally increasing the stimulus amplitude, the smallest displacement that elicited an AP with and without LSC activation was defined as the threshold for each condition. Mechanical stimuli were applied to corpuscles before and after experimental protocols to elicit mechanoreceptor APs, confirming health and proper function of the corpuscle and afferent throughout the experiment. All single-fiber and patch-clamp recordings were acquired from corpuscles in skin preparations from at least 3 different animals. Electrophysiological data was measured in Clampfit 10.7 (Molecular Devices), then analyzed and displayed in GraphPad Prism 9.5.1 (GraphPad Software, LLC).

### Enhanced Focused Ion Beam Scanning Electron Microscopy (eFIB-SEM)

eFIB-SEM procedures were performed as described previously (Nikolaev *et al*., 2023).

#### Sample preparation

A patch of bill skin was dissected from an E27 duck embryo and immediately immersed into fixative solution containing 2.5% glutaraldehyde, 2.5% paraformaldehyde, 0.13M cacodylate, 4 mM CaCl2, 4 mM MgCl2 (pH 7.4, 37°C). The epidermis was removed from the skin, which was then cut into 1 mm by 1 mm sections at room temperature. The dermis sections were then transferred to fresh fixative solution and gently shaken at 4°C for 48 hours. The solution was replaced with freshly prepared fixative solution at the 24-hour timepoint. After 48 hours, the sample was stored in a solution of 1.5% paraformaldehyde, 0.13M cacodylate pH 7.4 and stored at 4°C.

The bill skin samples were then sectioned into 300 µm thick slices in 0.13 M cacodylate buffer using a Compresstome (Precisionary, MA). The slices were washed in cacodylate buffer (0.13 M), postfixed with 2% osmium tetroxide and 1.5% potassium ferrocyanide in 0.13 M cacodylate buffer for 120 min at 0°C. After wash in distilled water, the slices were stained with 1% thiocarbohydrazide for 40 min at 40°C, 2% osmium tetroxide for 90 min at room temperature followed by 1% uranyl acetate at 4°C overnight. These staining reagents were diluted in the double distilled water. The sample slices were completely washed with distilled water between each step at room temperature three times for 10 min each. Finally, the slices were transferred into lead aspartate solution at 50°C for 120 min followed by distilled water wash at room temperature three times for 10 min each. After the heavy metal staining procedure, the samples were dehydrated with graded ethanol, embedded in Durcupan resin (Sigma, MO) and then polymerization at 60°C for 48 hours.

#### FIB-SEM sample preparation

Two duck bill skin samples embedded in Durcupan were selected for FIB-SEM sample preparation. The first sample, DB-01MP, included a Pacinian corpuscle in conjunction with a Meissner corpuscle, and the second sample from a different embryo, DB-02P, contained a larger Pacinian corpuscle. Each sample was first mounted on the top of a 1 mm copper post which was in contact with the metal-stained sample for belter charge dissipation, as previously described (Xu et al., 2017). Each vertical sample post was then trimmed to a small block with a width of 135 µm perpendicular to the ion beam, and a depth of 110 µm in the direction of the ion beam sequentially. Both blocks contain the Region of Interest (ROI) of one complete Pacinian corpuscle. The trimming for DB-01MP was guided by X-ray tomography data from a Zeiss Versa XRM-510, whereas DB-02P’s trimming used data from a Zeiss Versa XRM-620, both utilizing a Leica EM UC7 Ultramicrotome for trimming (Pang and Xu, 2023).

For conductive coating, a dual layer of 10-nm gold and 100-nm carbon was coated on the DB-01MP using a Gatan 682 High-Resolution Ion Beam Coater. The coating parameters were 6 keV, 200 nA on both argon gas plasma sources, 10 rpm sample rotation with 45-degree tilt. Conversely, DB-02P was first coated with 10-nm gold using a Cressington Sputter Coater 208HR, rotating at 15 rpm with a 30-degree tilt using 40 mA argon plasma source, followed by a 40-nm carbon layer deposited using a Leica ACE200 carbon coater.

#### FIB-SEM 3D large volume imaging

Two FIB-SEM prepared samples, DB-01MP and DB-02P were imaged using two enhanced FIB-SEM systems, as previously described (Xu *et al*., 2017; Xu et al., 2020; Xu et al., 2021). For DB-01MP, the ROI block face was imaged with a 2 nA electron beam at 2 MHz scanning rate and a landing energy of 1.2 keV, while for DB-02P, it was scanned with a 3 nA electron beam at 3 MHz under the same landing energy condition. Both samples had an x-y pixel size set at 8 nm. A subsequently applied focused Ga^+^ beam of 15 nA at 30 keV strafed across the top surface and ablated away 8 nm of the surface. The newly exposed surface was then imaged again. The ablation – imaging cycle continued about once every minute for one week to complete DB-01MP that contains one Meissner and one Pacinian corpuscle, and about once every minute for two weeks to complete DB-01P that contains a larger Pacinian corpuscle from a different embryo. The acquired image stack formed a raw imaged volume, followed by post processing of image registration and alignment using a Scale Invariant Feature Transform (SIFT) based algorithm. The aligned stack consists of a final isotropic volume of 85 x 56 x 75 µm^3^ and 94 x 86 x 120 µm^3^ and for DB-01MP and DB-02P, respectively. The voxel size of 8 x 8 x 8 nm^3^ was maintained for both samples throughout entire volumes, which can be viewed in any arbitrary orientations.

#### Electron microscopy segmentation

The segmentation of organelles, cells, and subcellular structures from EM images was achieved with ZEISS arivis Cloud, an AI-driven cloud-based platform (https://www.apeer.com/) (Dang et al., 2021). Deep learning techniques were utilized to achieve automated segmentation, employing a customized convolutional neural network (CNN) architecture based on 2D U-Net. To generate ground truth data, cells and organelles were manually annotated from a small set (100 planes) of the raw EM images. The CNNs were trained using the annotated ground truth data and proofread to achieve high-quality segmentation of the objects in 3D. Semantic segmentation was applied to each object, and the accuracy of the segmentation was assessed by evaluating the voxel Intersection over Union (IoU) and F1 scores. IoU was calculated as the overlap between annotation and ground truth bounding boxes by computing the ratio of the intersection area to the union area: IoU = (Intersection Area) / (Union Area). The F1 score was calculated as the balance between the model’s ability to correctly identify positive samples (precision) and its ability to capture all positive samples (recall): F1 = 2 * (Precision * Recall) / (Precision + Recall) (Padilla et al., 2020). Arivis machine learning models were downloaded separately for each class of cells or organelles to create a full 3D model on a full dataset. All volumes were segmented at 8 x 8 x 8 nm.

#### Electron microscopy reconstruction and data analysis

Raw EM data, along with ZEISS arivis machine learning models for each class, were imported into ZEISS Arivis Pro software. This software was used to segment each individual cell and organelle, creating complete objects. In certain cases, ZEISS arivis Hub from the FIB-SEM Collaboration Core was used to generate the objects. These objects were then filtered by size to eliminate any extraneous noise components. Manual proofreading and adjustments were made as necessary. Various quantitative measures, including volume, distances, surface area, and diameters, were calculated within the software. Videos were generated using Arivis Pro. The 3D TEM tomography was reconstructed at a resolution of 1.6 x 1.6 x 1.6 nm.

### 3D transmission electron microscopy tomography

Procedures were performed as described previously (Nikolaev *et al*., 2023). Freshly peeled duck bill skin was fixed in Karnovsky fixative at 4°C for 1 hour, washed in 0.1 M sodium cacodylate buffer (pH 7.4), then postfixed in 1% osmium tetroxide for 1 hour in the dark on ice. The tissue was stained in Kellenberger solution for 1 hour at room temperature after washing in distilled water, dehydrated in a series of alcohols and propylene oxide, then embedded in EMbed 812, and polymerized overnight at 60°C. Thick sections of 250 nm depth were obtained from hardened blocks using a Leica UltraCut UC7 on copper formvar coated slot grids. 250 nm thick sections were contrast stained using 2% uranyl acetate and lead citrate and 15nm fiducial gold was added to both sides to aid alignment for Tomography. Sections were viewed using a FEI Tecnai TF20 at 200 Kv and data was collected using SerialEM (Mastronarde, 2005) at voxel size of 1.6 x 1.6 x 1.6 nm^3^ on a FEI Eagle 4Kx4K CCD camera using tilt angles of -60 to 60 degrees. All solutions were supplied by Electron Microscopy Sciences (Hatfield, PA).

### RNA sequencing of Pacinian inner cores

Single inner cores of Pacinian corpuscles were manually isolated and collected from the *ex vivo* bill-skin preparation in RNase-free conditions for eventual transcriptomic analysis. First, the inner core was separated from the outer core using Krebs-filled patch pipettes with large pressure applied via a High-Speed Pressure Clamp, as described above for electrophysiology. Aspiration pipettes with tip diameters of ∼50–100 µm filled with 3 µl of RNA Lysis Buffer (Zymo) were then used to aspirate the inner core by applying light negative pressure. The lysis buffer with the inner core from the pipette was then deposited into a 1.5 ml microcentrifuge tube using positive pressure and 10 µl of extra RNA Lysis Buffer was added to each tube. Samples were then stored at −80°C until RNA isolation. RNA was isolated using the Quick-RNA Microprep Kit (Zymo) per the manufacturer’s instructions. RNA concentrations of 113–782 pg/µl and RIN values in the range of 7.1–9.5 were acquired from inner cores, assessed via a 2100 Bioanalyzer (Agilent). RNA from a total of 6-7 inner cores was collected from 5 independent embryos. RNA was also isolated from 6 epidermis samples of 5 separate bill-skin preparations as a control.

Library preparation and sequencing were performed by the Yale Center for Genome Analysis. Libraries were prepared using the NEBNext Single Cell/Low Input RNA Library Prep Kit (New England Biolabs) and sequencing was done with an Illumina NovaSeq instrument in the 100 bp paired-end mode. Approximately 35-87 million sequencing read pairs per sample were obtained. The raw sequencing data was subsequently processed on the Yale Center for Research Computing cluster. First, raw reads were filtered and trimmed using Trimmomatic v0.39 with default parameters. Filtered high-quality reads were then aligned to the duck reference genome using the STAR aligner v2.7.9a with default parameters.

The duck reference genome

ftp://ftp.ncbi.nlm.nih.gov/genomes/all/GCF/000/355/885/GCF_000355885.1_BGI_duck_1.0/GCF_000355885.1_BGI_duck_1.0_genomic.fna.gz

and gene annotation

ftp://ftp.ncbi.nlm.nih.gov/genomes/all/GCF/000/355/885/GCF_000355885.1_BGI_duck_1.0/GCF_000355885.1_BGI_duck_1.0_genomic.gff.gz

were obtained from the National Center for Biotechnology Information. The gene annotation was filtered to include only protein-coding genes. Aligned reads were counted by the featureCounts program within the Subread package v2.0.1 with default parameters. FPKM values were calculated from read counts using the edgeR v3.34.1 package (Bioconductor v3.13) in R v4.1. Statistical analysis of differential expression of genes between groups was evaluated using the Fisher’s Exact Test with the Benjamini-Hochberg method for false discovery in edgeR. RNA sequencing data were deposited to the Gene Expression Omnibus, accession number GSE273272.

To compare the transcriptomic signature of the Pacinian inner core with that of Meissner corpuscles, previously published transcriptomic data from Meissner corpuscles was used (Nikolaev *et al*., 2023). Transcriptomic data from Pacinian inner core, epidermis, Meissner corpuscles, and bill skin (dermis) was jointly reanalyzed according to the same pipeline described above. Principial component analysis was performed on log-transformed normalized (counts per million) expression data using *prcomp* function in R. First two principal components were extracted. Group means were determined for the first two principal components. Euclidean distance based on first two principal components between group means was determined using *dist* function in R.

## Resource availability

### Lead contact

Requests for further information and resources should be directed to the lead contact, Sviatoslav N. Bagriantsev.

### Materials availability

This study did not generate new unique reagents.

### Data and code availability

All data are available in the main text or the supplementary materials. RNA sequencing data are deposited to the Gene Expression Omnibus, accession number GSE273272. This study does not report original code.

## Supplementary Information

**Figure S1.**
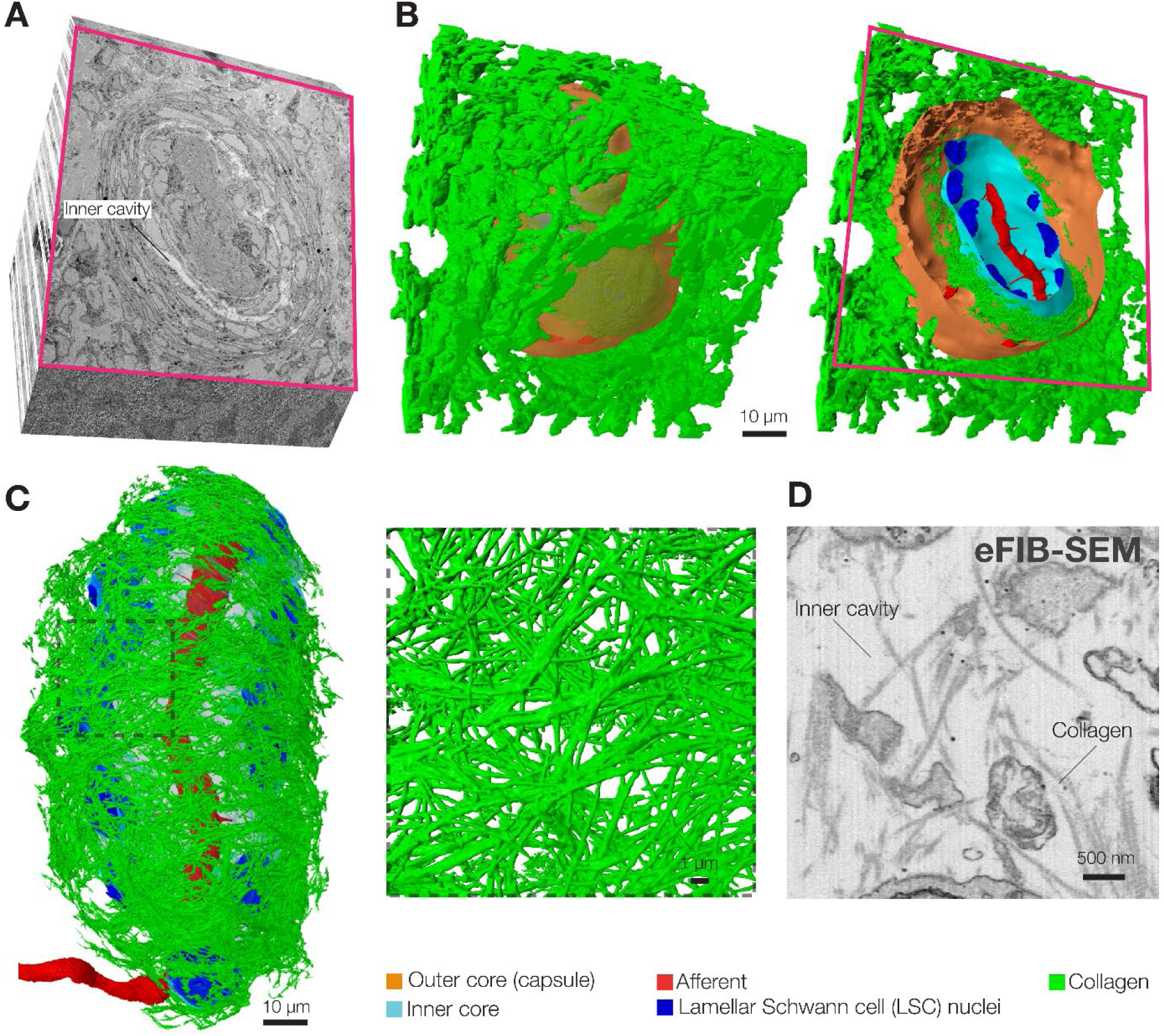
The cavity between the outer and inner cores is filled with collagen fibers. (A) A cross-section of the 3D volume from eFIB-SEM data showing the location of the inner cavity in the Pacinian corpuscle. (B) 3D reconstruction of the same volume and a cross-section, as in (A), showing thick collagen bundles surrounding the outer core. (C) A 3D reconstruction of a Pacinian inner core surrounded by single collagen fibers, along with a magnified region. (D) eFIB-SEM single image from the inner cavity showing single collagen fibers inside the corpuscle.

**Figure S2.**
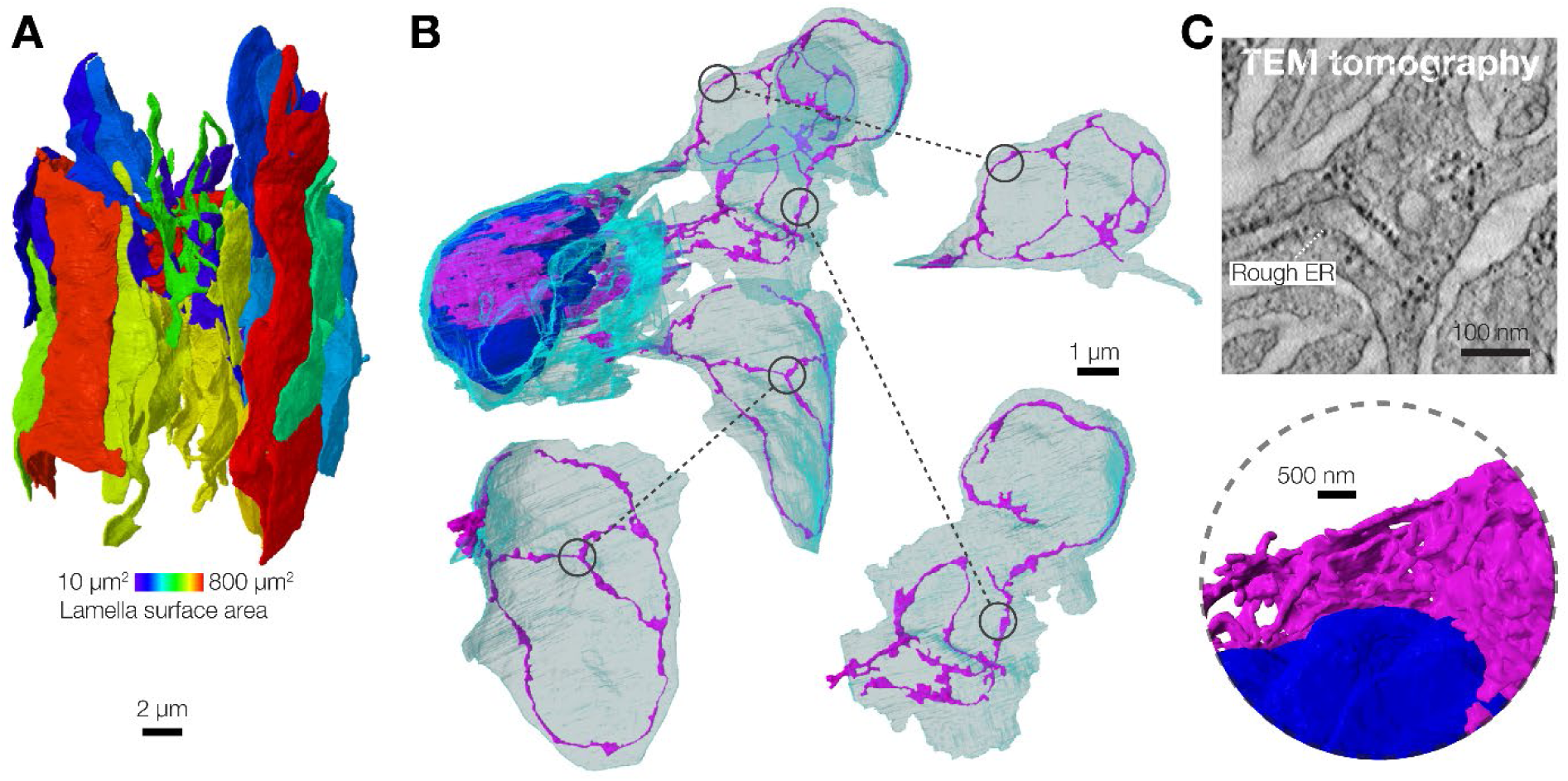
3D reconstruction of a lamellar Schwann cell from the Pacinian inner core. (A) 3D reconstruction of an LSC showing surface area of lamellae. (B, C) 3D reconstruction of endoplasmic reticulum (B) and a single FIB-SEM image of rough endoplasmic reticulum (rough ER) in LSC lamellae (C).

**Figure S3.**
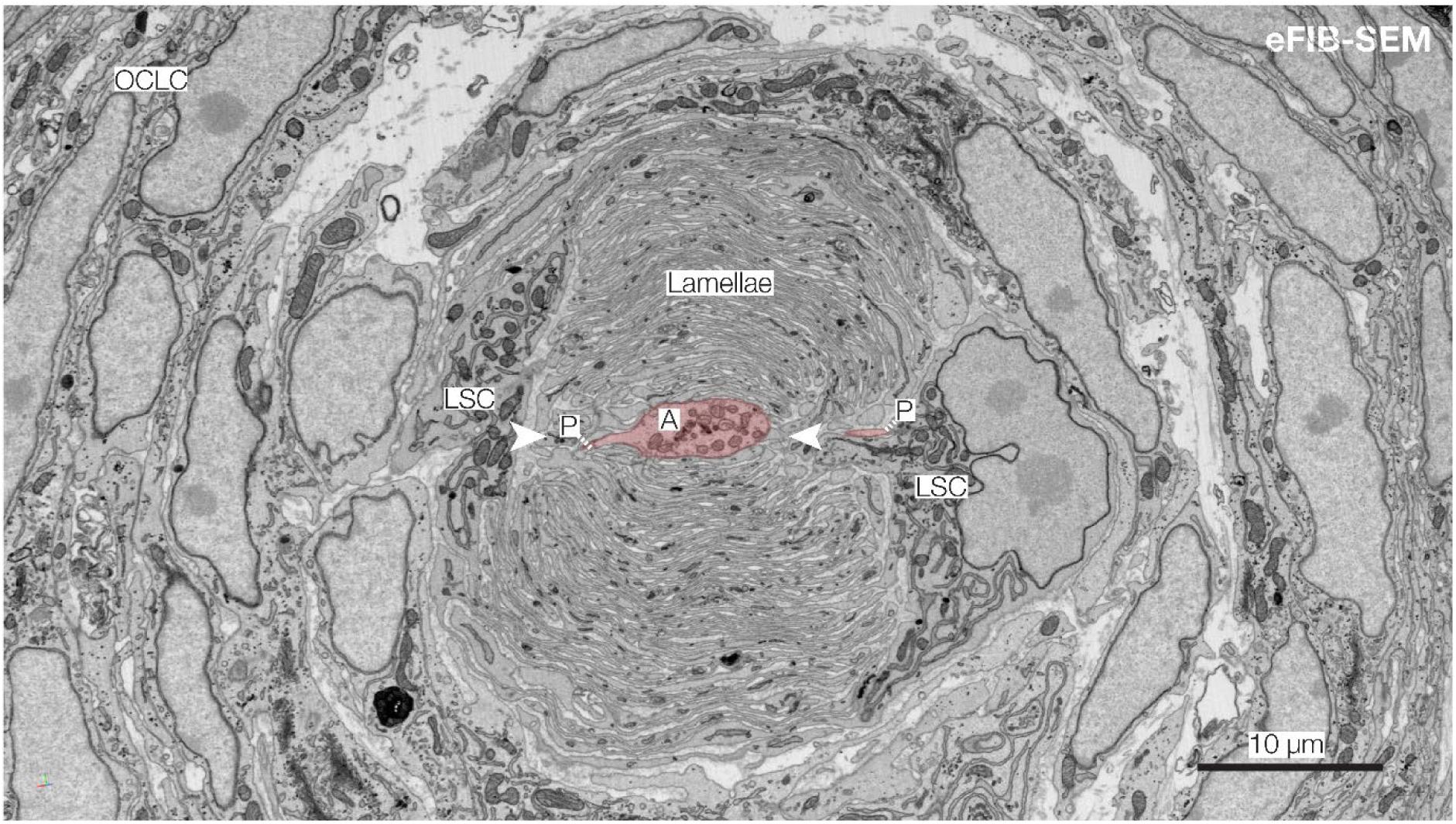
Protrusions emanate from the narrow sides of afferent terminal facing the cleft in LSC lamellae. Shown is an eFIB-SEM image of a Pacinian inner core. LSC, lamellar Schwann cells, OCLC, outer core lamellar cells, N, LSC nucleus; A, afferent terminal ; P, protrusion. Arrowheadspoint to the cleft formed by inner core lamellae.

**Figure S4.**
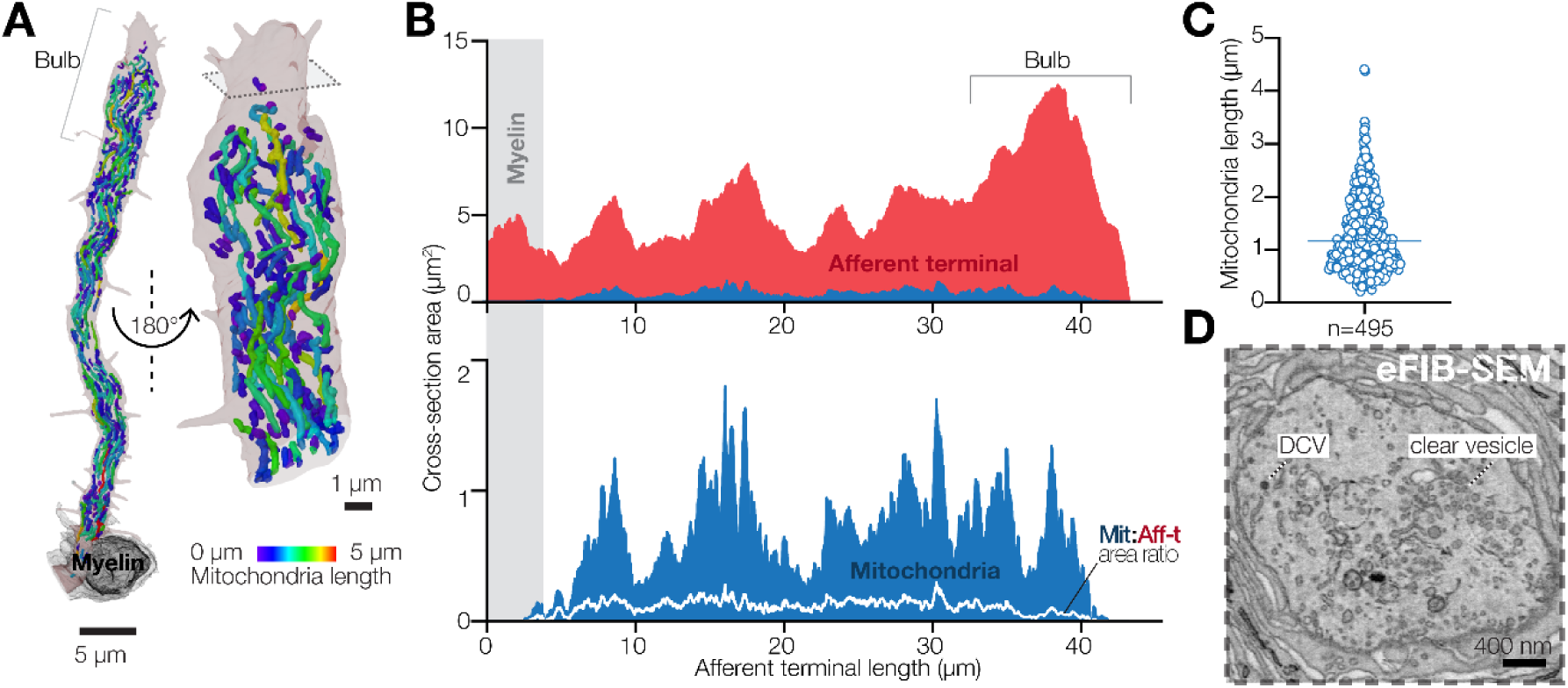
Mitochondria and vesicles in the Pacinian afferent terminal. (A) 3D reconstruction of mitochondria in the afferent terminal. (B) Quantification of cellular area occupied by mitochondria along the length of the terminal. (C) Quantification of mitochondria length in the afferent terminal. (D) A single eFIB-SEM image of the afferent terminal bulb with clear vesicles and dense core vesicles (DCV).

**Figure S5.**
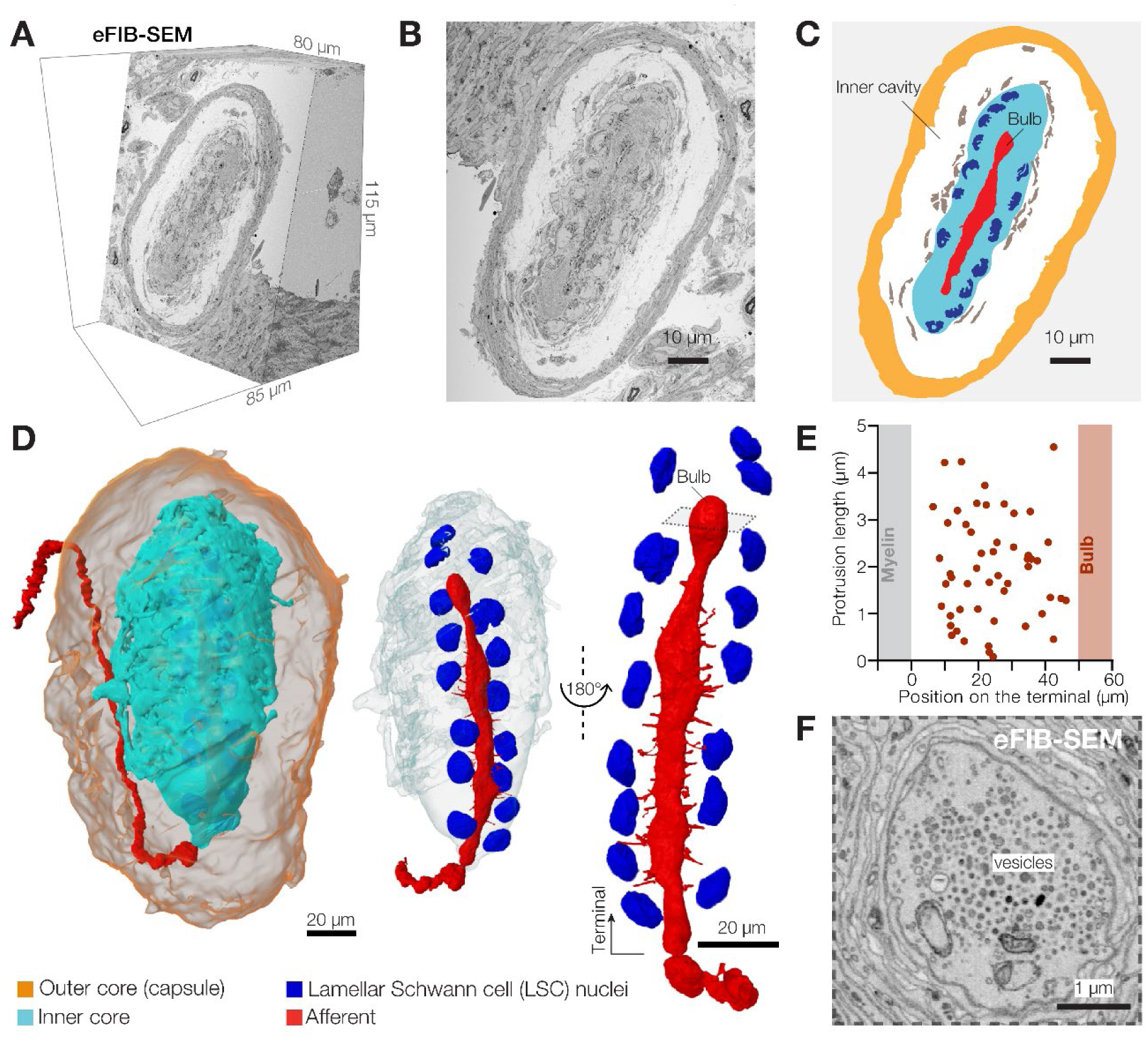
3D architecture of a second Pacinian corpuscle. (A) A 3D volume of duck bill skin dermis obtained by eFIB-SEM. (B, C) A single eFIB-SEM image (B) and an illustration (C) of a section of the Pacinian corpuscle. (D) 3D reconstruction of the Pacinian corpuscle showing the location of the inner core inside the outer core, and reconstruction of the inner core and the afferent. (E) Localization, length and target of afferent protrusions in the second Pacinian. (F) A single eFIB-SEM image of the bulb area of the afferent terminal showing clear and dense core vesicles (DCV).

**Figure S6.**
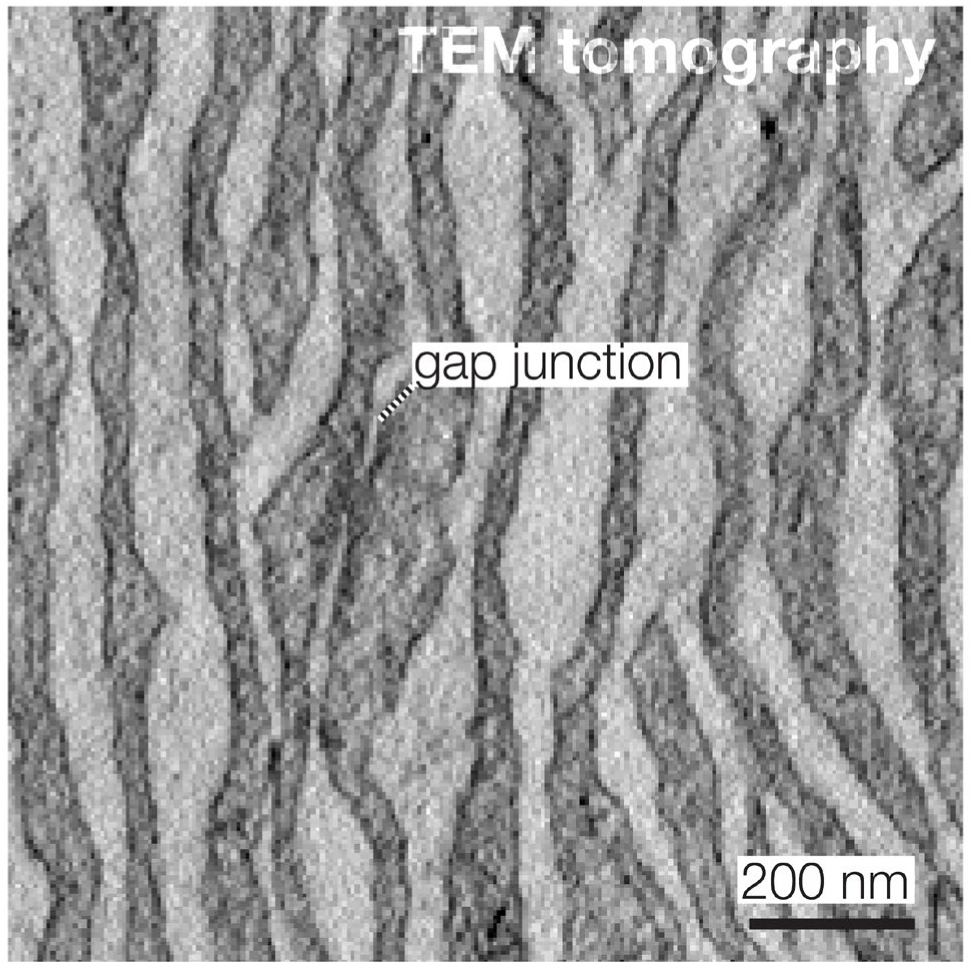
LSC lamellae are connected by gap junctions. Shown is a transmission electron microscopy image of inner core lamellae connected by a gap junction.

**Figure S7.**
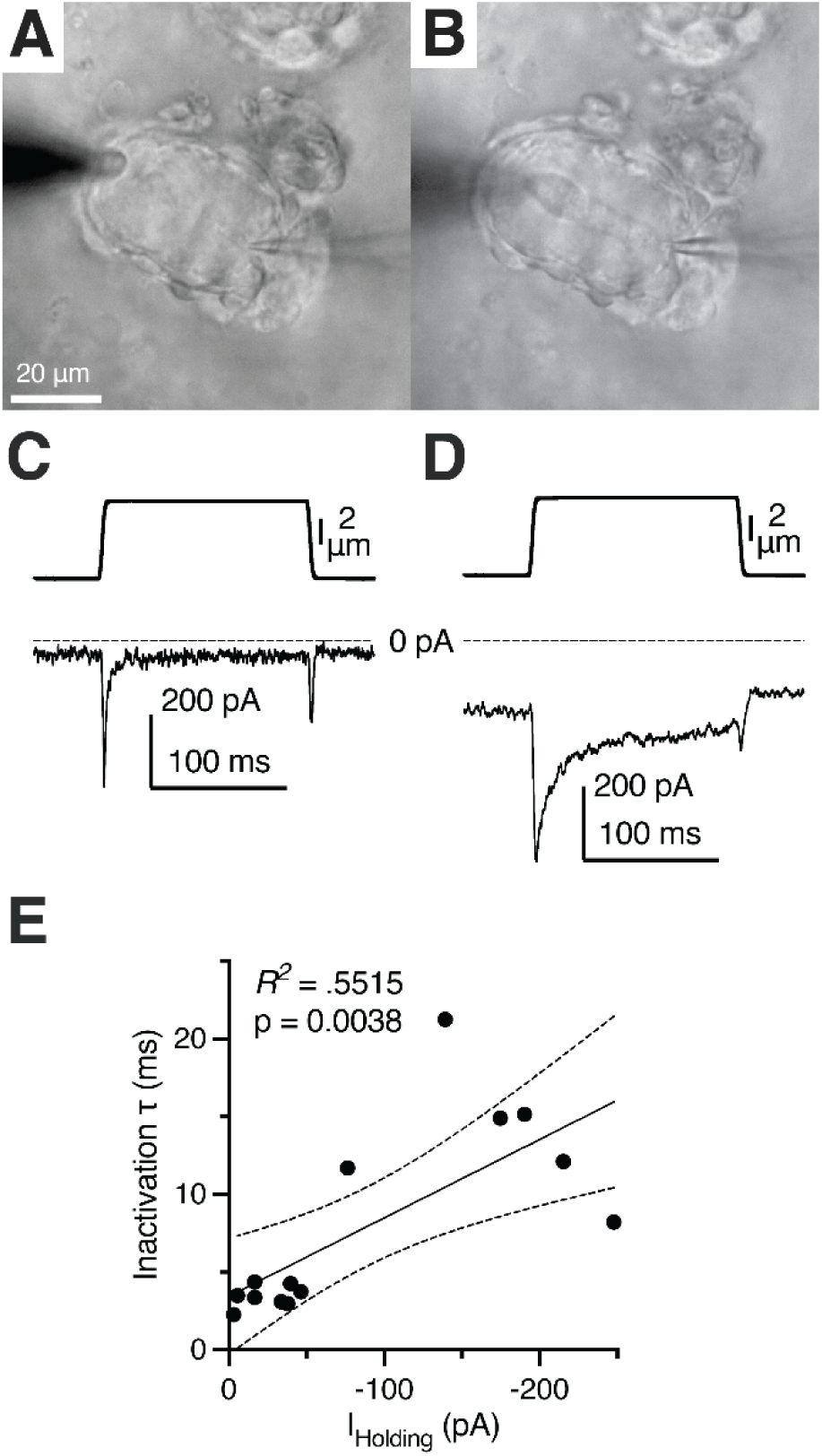
Prolongation of MA current inactivation in a deteriorated Pacinian afferent terminal after decapsulation. (A) Bright-field image of a patched Pacinian terminal in a decapsulated corpuscle. (B) Bright-field image of the deteriorated decapsulated terminal from (A). (C) Mechanical stimulus (top) and MA current response (bottom) of the patched terminal in (A) after decapsulation shows fast kinetics of inactivation. (D) Mechanical stimulus (top) and MA current response (bottom) of the deteriorated decapsulated terminal at the same time point that the image in (B) was captured. (E) Relationship between inactivation rate of MA current in patch-clamped terminals and the holding current applied during voltage clamp at -60 mV, fitted to the linear equation.

**Figure S8.**
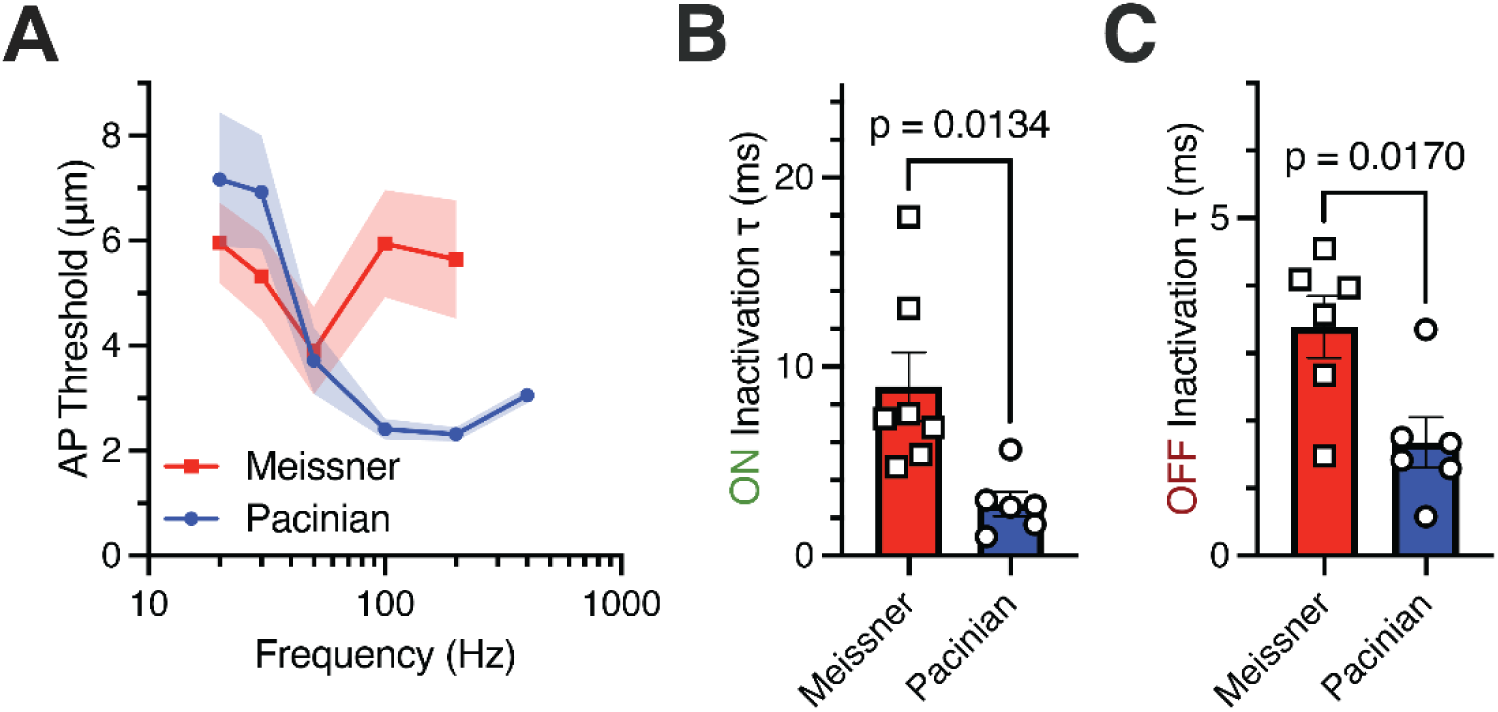
Correlation between peak frequency sensitivity and MA current inactivation in Meissner and Pacinian corpuscles. (A) Population tuning curves recorded from afferents of avian Meissner and Pacinian corpuscles. Data are shown as mean ± SE from 9 Meissner and 26 Pacinian corpuscles. (B, C) Rates of MA current inactivation (B, ON response; C, OFF response) recorded from Meissner and Pacinian afferent terminals. Symbols represent recordings from individual corpuscles. Data are shown as mean ± SEM. Statistics: Welch’s t-test.

**Figure S9.**
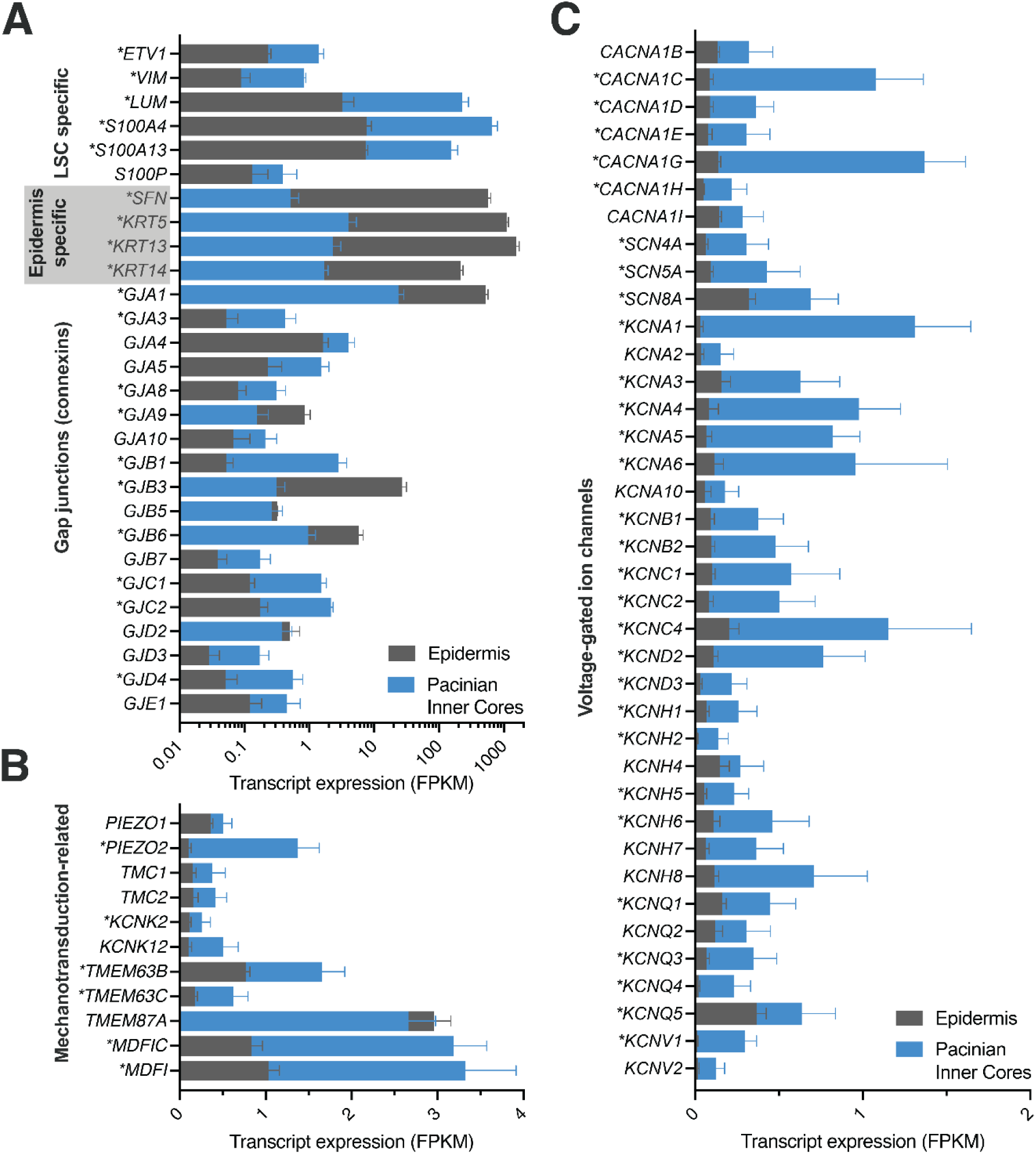
RNA sequencing of Pacinian inner cores. (A-C) Shown are fragments per kilobase per million reads sequenced from mRNA of genes of (A) gap junction connexins, (B) known and putative mechanically-gated ion channels and their modifiers, (C) voltage-gated sodium, voltage-gated calcium channels, and voltage-gated potassium channels from isolated Pacinian inner cores, compared to expression of such genes in the duck bill epidermis. Data are mean + SEM from 7 inner cores and 6 epidermis samples. Statistics: Fisher’s Exact Test with Benjamini-Hochberg method for false discovery rate (FDR) *FDR-adjusted P < 0.05.

**Figure S10.**
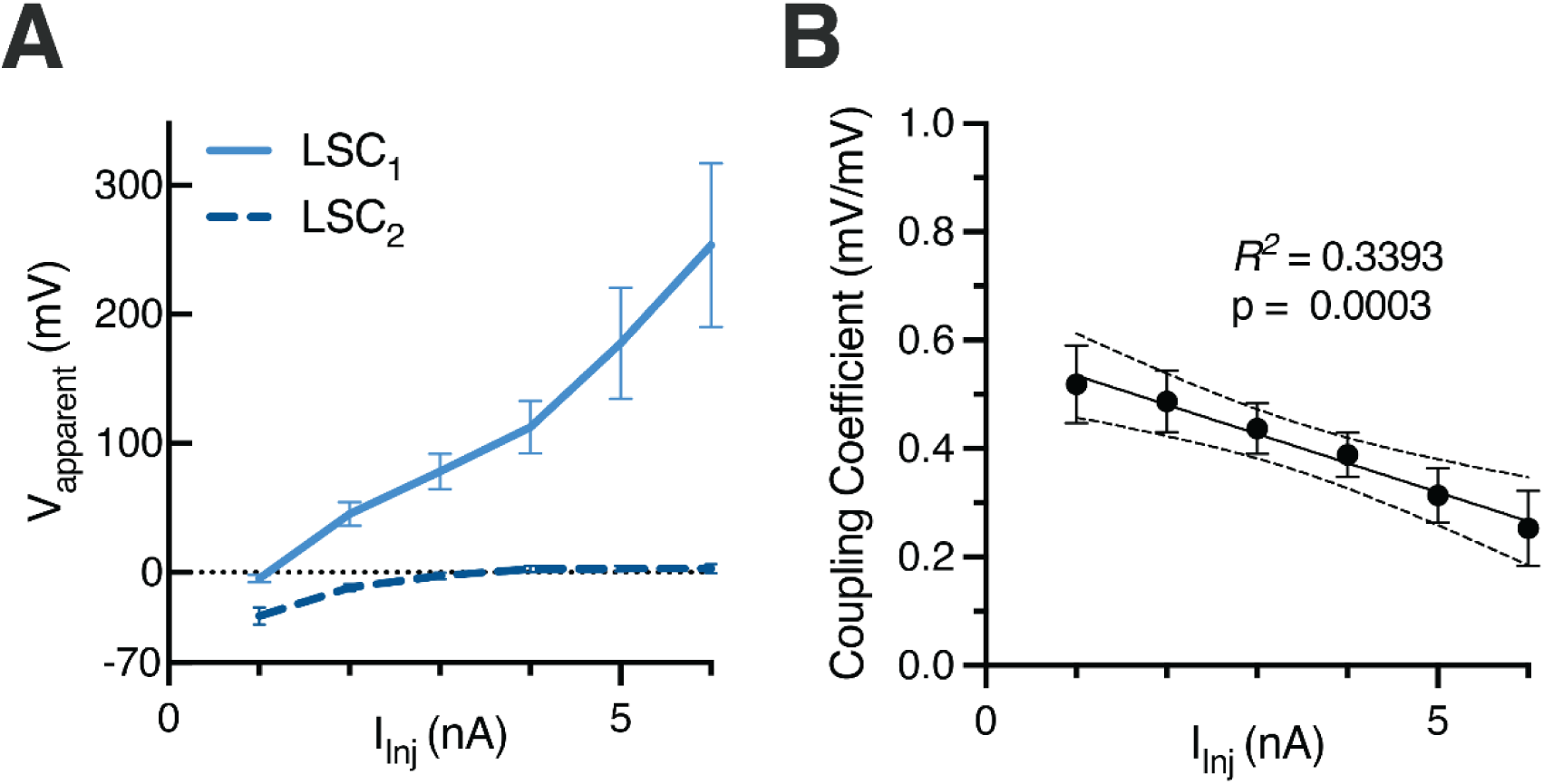
Electrical coupling between adjacent LSCs is voltage-dependent. (A) Quantification of the effect of current injection into an LSC on the apparent membrane potential of the same and adjacent LSC (LSC_1_ and LSC_2_, respectively). Data are mean ± SEM from 5 recordings. (B) The coupling coefficient between LSC pairs from (A). Data are mean ± SEM from 5 recordings, fitted to the linear equation.

**Figure S11.**
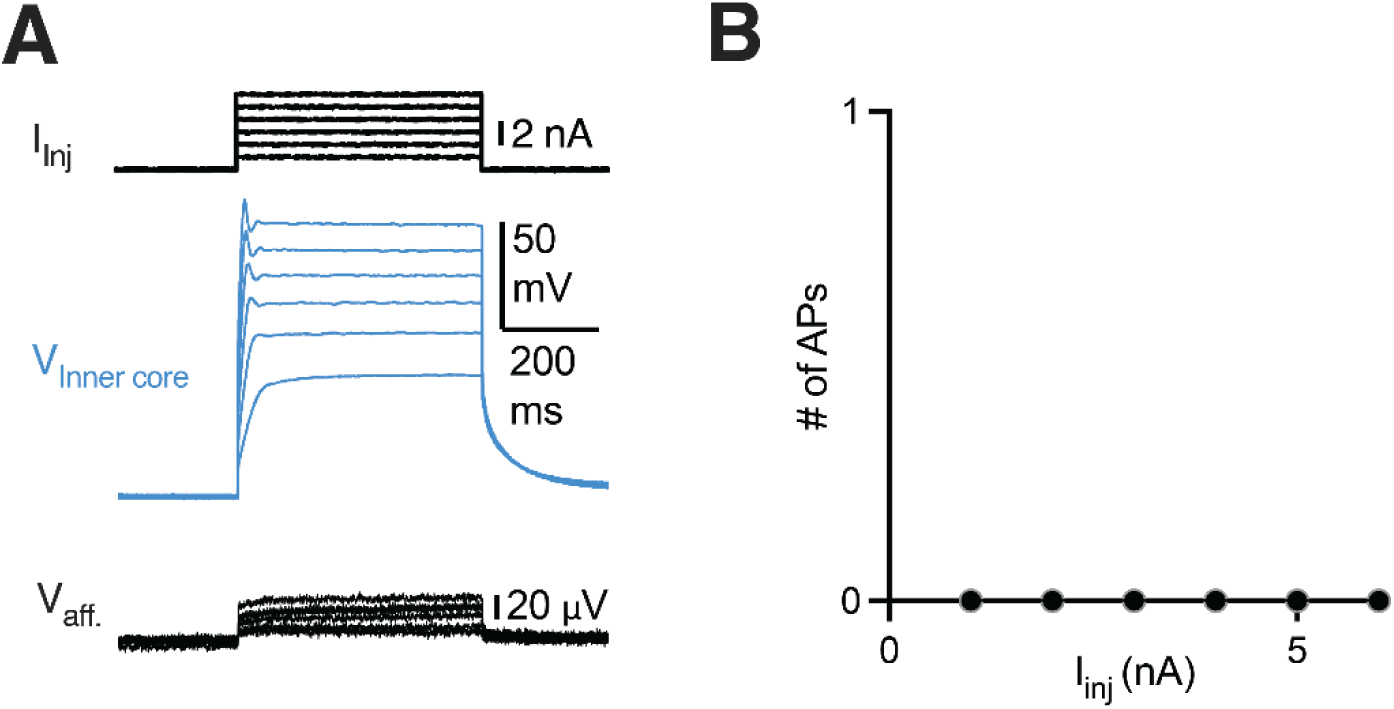
Activation of a single LSC by current injection fails to induce AP firing in the afferent. (A) Current injection stimulus applied to a patch-clamped LSC (top), voltage response of the patched LSC (middle), and single-fiber response of the associated Pacinian afferent during simultaneous paired recording. (B) Quantification of the number of action potentials elicited during LSC activation by current injection. Data shown as overlapping lines representing individual cells from 6 recordings (all 0 APs).

**Fig S12.**
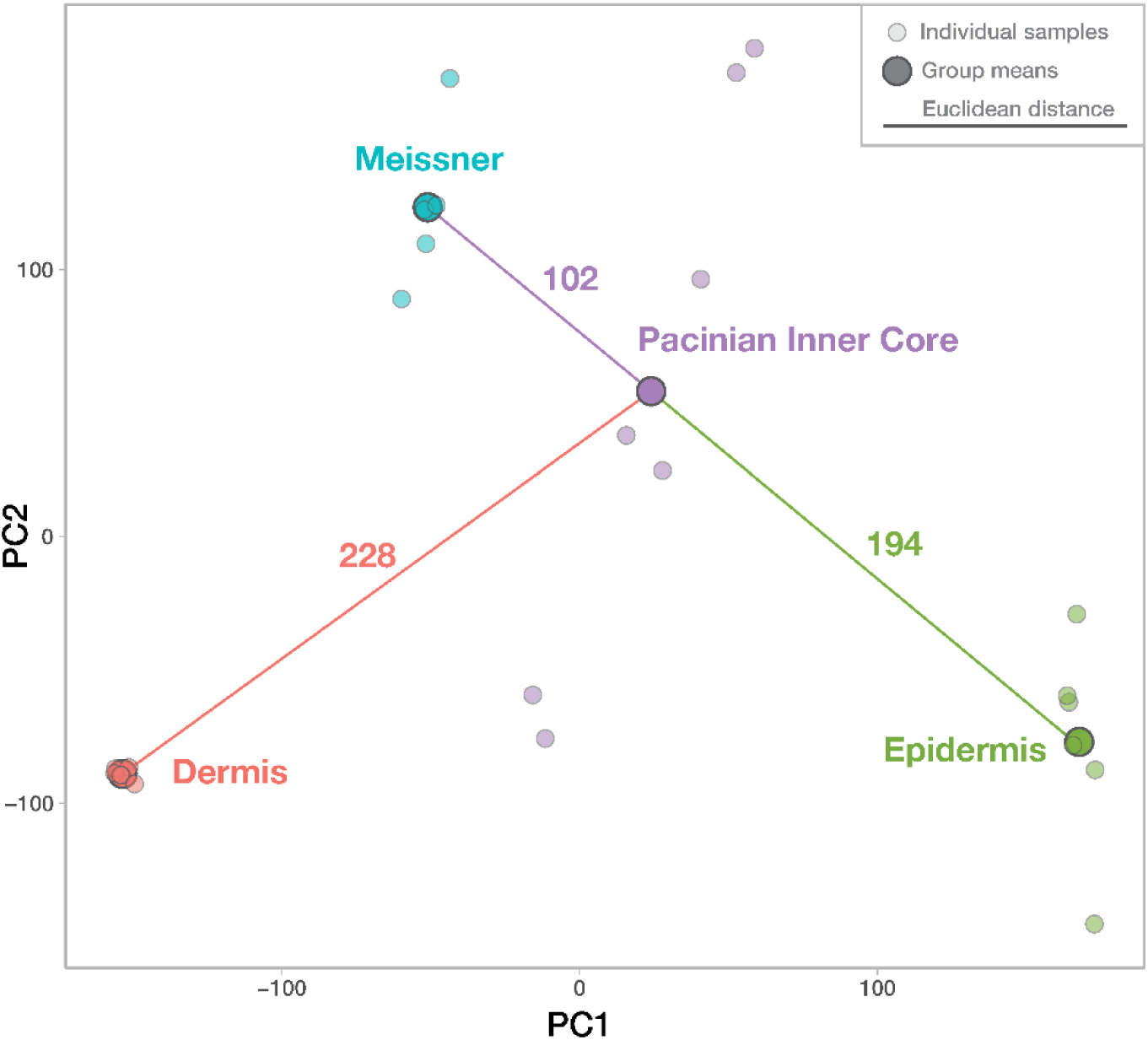
Pacinian inner core is transcriptomically more similar to Meissner corpuscles than to bill skin epidermis or dermis. A PCA plot (first two principal components) of transcriptomic data from Pacinian inner core and bill skin epidermis (this study), Meissner corpuscles and bill skin dermis (Nikolaev *et al*., 2023) showing individual samples (small circles) and group means (large circles). Numbers above the connecting lines indicate Euclidean distances between group means. Activation of a single LSC by current injection fails to induce AP firing in the afferent.

**Table S1.** Accuracy statistics of eFIB-SEM data segmentation for Pacinian corpuscles.

| Method | Object | IoU* | F1 score |
| --- | --- | --- | --- |
| eFIB-SEM Pacinian #1 | Afferent1 | 0.67 | 0.80 |
|  | Inner core | 0.76 | 0.86 |
|  | Outer core | 0.55 | 0.71 |
|  | Collagen | 0.65 | 0.79 |
|  | ER (lamellae) | 0.75 | 0.86 |
|  | ER (cell body) | 0.85 | 0.92 |
|  | Nuclei | 0.78 | 0.88 |
|  | Mitochondria | 0.58 | 0.73 |
|  | Lamellae | 0.75 | 0.86 |
| eFIB-SEM Pacinian #2 | Afferent2 | 0.85 | 0.92 |
| TEM tomography Pacinian #1 | Afferent | 0.98 | 0.99 |
|  | Lamellae | 0.95 | 0.97 |
|  | Tethers | 0.78 | 0.88 |

\*Intersection over Union

**Movie S1.** 3D architecture of an avian Pacinian corpuscle obtained using eFIB-SEM.

**Movie S2.** 3D reconstruction of a fragment of lamellar cell-afferent contact area obtained by transmission electron microscopy tomography.

**Supplementary Data S1.** RNA sequencing of Pacinian inner cores vs epidermis. Data were deposited to the Gene Expression Omnibus, accession number GSE273272.

